# Restoring Shugoshin 1 reduces chromosome errors in human eggs

**DOI:** 10.64898/2026.01.08.698387

**Authors:** Debojit Saha, Saba Manshaei, Tommaso Cavazza, Zuzana Holubcová, Barbora Maierova, Agata P. Zielinska, Lena Wartosch, Martyn Blaney, Kay Elder, Melina Schuh

## Abstract

Aneuploidy in human eggs, which rises sharply with age, is a leading cause of infertility, IVF failure, and miscarriage. This age-related aneuploidy is primarily driven by premature sister chromatid separation (PSSC), resulting from loss of the cohesin complex that holds chromatids together. How cohesin is destabilized in the long-lived mammalian oocyte is poorly understood. Here, we show that in mouse oocytes, pericentromeric transcription is essential for maintaining the cohesion protector Shugoshin 1 (SGO1) and PP2A at centromeres, which together safeguard the cohesin subunit REC8. With age, mouse oocytes lose pericentromeric transcription, SGO1, and PP2A, leading to destabilized cohesion and increased PSSC. Supplementing aged mouse oocytes with *Sgo1* restores centromeric protection, and reduces PSSC to youthful levels. Aged human oocytes also show reduced SGO1, and *SGO1* supplementation reduces the fraction of human eggs with PSSC by approximately half. These findings establish *SGO1* supplementation as a potential strategy to preserve chromatid cohesion in aging oocytes.

## Introduction

Aneuploidy in human eggs results from errors in chromosome segregation during meiosis and is the leading cause of infertility, in vitro fertilization (IVF) failure, miscarriage, and congenital disorders in the offspring of women of advanced maternal age^1–3^. The molecular mechanisms underlying these segregation errors remain incompletely understood^4–6^.

Meiosis in oocytes is a prolonged process that spans decades in humans. In female mammals, oocytes initiate meiosis during fetal development and arrest at the germinal vesicle (GV) stage in prophase I. Upon hormonal stimulation, they resume meiosis, complete meiosis I (MI) to segregate homologous chromosomes, and arrest again at metaphase II (MII), where sister chromatids remain linked until fertilization.

Sister chromatids should remain linked during meiosis I and separate only during meiosis II, following fertilization. However, they sometimes separate prematurely during MI and MII. Because the resulting single chromatids can no longer be correctly aligned on the spindle, they distribute randomly during anaphase, increasing the risk of aneuploidy. This premature separation of sister chromatids (PSSC) arises from defects in chromosome cohesion and is the leading cause of age-related aneuploidy^7–11^.

Chromosome cohesion and accurate chromosome segregation depend on the cohesin complex, which holds sister chromatids together and is loaded during fetal development^12–14^. Cohesin is thought not to be replenished in mammalian oocytes, unlike somatic cells^15–17^. Thus, oocytes are especially vulnerable to cohesion loss as females age. While the gradual loss of cohesin with age in prophase-arrested oocytes is well documented, the molecular events that underlie its destabilization and trigger PSSC upon meiotic resumption are poorly defined^18–22^.

Shugoshin proteins protect centromeric cohesin by recruiting the phosphatase PP2A, which counteracts the phosphorylation-dependent cleavage of the cohesin complex subunit REC8 by the enzyme Separase^23–26^. SGO2 is reported to be essential for centromeric cohesion protection and proper chromosome segregation in mammalian oocytes, however the role of its paralog, SGO1, is unresolved^27–34^.

In mitotic cells, SGO1 localization and centromeric cohesion are regulated not only by structural chromatin features, but also by dynamic processes involving BUB1 kinase, histone phosphorylation, and RNA polymerase II-dependent transcription^35–43^. Transcription at centromeres, once considered transcriptionally inert, plays a critical role in recruiting SGO1, maintaining centromeric cohesion, and enabling proper kinetochore function. Specifically, transcription and noncoding RNAs produced at centromeres are proposed to release SGO1 from kinetochores, which sit on the outer face of each sister centromere. This release allows SGO1 to move to the inner centromere, the region between sister chromatids, where it can engage with cohesin^44,45^. Although centromeric noncoding RNAs have been implicated in kinetochore function and cohesion maintenance during mitosis, their role in meiosis remains incompletely understood^46–48^. It is also unclear whether a transcription-dependent mechanism for cohesion protection operates in oocytes or is compromised with maternal age. These gaps limit our ability to understand how cohesion is preserved in female meiosis and to identify strategies to prevent age-related aneuploidy.

Aneuploidy in human eggs remains a major barrier in reproductive medicine, and no interventions currently exist to preserve chromosome cohesion in human eggs^49–54^. To address this unmet need, we investigate the roles of transcription and SGO1 in cohesion in mouse and human eggs and whether their decline with age contributes to PSSC^55–57^. Our findings uncover a previously unrecognized mechanism of cohesion deprotection in aging oocytes and suggest a therapeutic strategy for improving human egg quality and IVF success rates.

## Results

### SGO1 prevents PSSC in mouse eggs

The role of SGO1 in meiotic cohesion has remained unresolved due to conflicting reports^28,29,33,58^. To address these discrepancies, we examined the subcellular localization SGO1 by microinjecting mouse GV oocytes with mRNAs encoding fluorescently labeled SGO1 and by detecting endogenous SGO1 by immunofluorescence. SGO1 localized to an extended region spanning the centromeres and pericentromeres at both metaphase I and metaphase II (Figures 1A–C). To directly test the importance of SGO1 in oocyte cohesion, we depleted endogenous SGO1 using two independent siRNAs in follicle-enclosed oocytes cultured for 10 days^59^. Following culture, oocytes were matured in vitro and analyzed at the MII stage. Both siRNAs led to a marked reduction of SGO1 on chromosomes of MII eggs (Figures 1C and 1D). Importantly, depletion of SGO1 resulted in a significant increase in premature sister chromatid separation (PSSC) at metaphase II compared to controls (Figures 1C–E). Prematurely dissociated sister chromatids cannot stably align at the spindle equator. As expected, SGO1-depleted eggs showed a significant increase in alignment defects (Figures 1C and 1F). We observed a higher fraction of oocytes with chromosome errors compared with previous studies using shorter depletion protocols^29,58^. This likely reflects more complete SGO1 loss after prolonged culture. We infer that SGO1 is essential for maintaining chromosome integrity and preventing PSSC in mouse oocytes.

**Figure 1.**
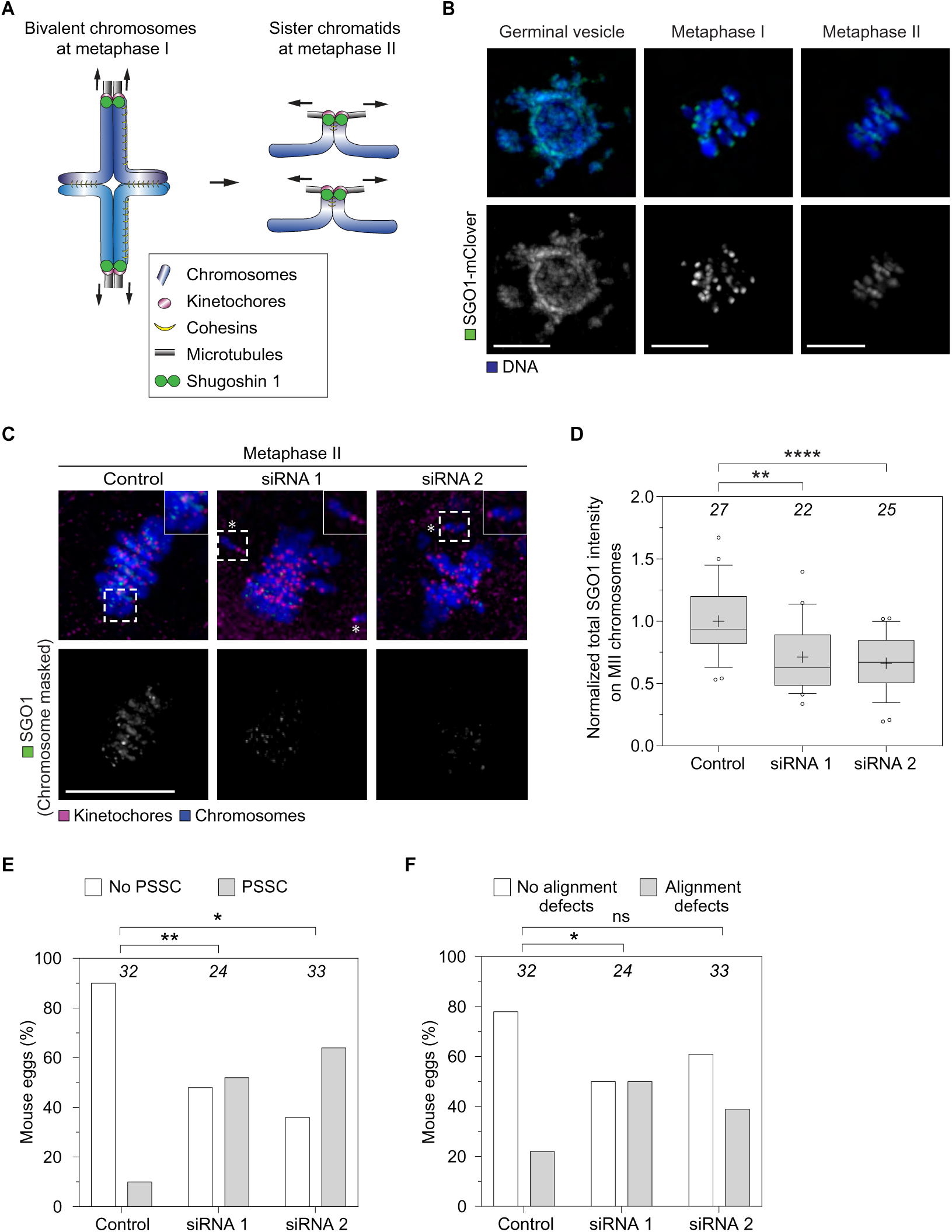
Shugoshin 1 protects sister chromatid cohesion in mouse eggs. (A) Schematic representation of the localization of the cohesin complex and the centromeric cohesin protector SGO1 in mouse oocytes at MI (left panel) and eggs at MII (right panel). (B) Representative time-lapse images of mouse oocytes at GV and MI stage, and eggs at MII stage microinjected with mRNA encoding mClover-SGO1 (green). DNA (H2B-SNAP) is shown in blue. Scale bars: 10 µm. (C–F) Loss of SGO1 from early-stage eggs results in increased PSSC. Representative images of metaphase II chromosomes (in blue) from mouse eggs treated with negative control (left panel), Sgo1 siRNA 1 (middle panel) and Sgo1 siRNA 2 (right panel) show that SGO1 (in green, chromosome masked) is lost upon knockdown. White asterisks indicate the presence of single chromatids. Kinetochores (CREST, magenta), DNA (Hoechst, blue) are shown in the top panel. Scale bar: 10 µm. (D) Mean fluorescence intensity of SGO1 on chromosomes of MII eggs after SGO1 knockdown. Plots show n = number of eggs analyzed at the top. mean (plus), median (horizontal black line), 25th and 75th percentiles (boxes), 10th and 90th percentiles (whiskers), and the 1st and 99th percentiles (circles). Statistical significance was measured by the Mann-Whitney test, **p≤0.01, ****p≤0.0001. (E–F) The percentage of eggs with PSSC (E) and chromosome alignment defects (F) increases after SGO1 knockdown with siRNA 1 and siRNA 2. Plots show n = number of eggs analyzed at top. Statistical significance was measured by Fisher’s exact test on the number of eggs with or without PSSC. ns = not significant, *p≤0.05, **p≤0.01.

### Transcription is required to maintain SGO1 and PP2A and prevent PSSC in oocytes

Centromeric transcription is reported to facilitate SGO1 recruitment during mitosis in tissue culture cells^36,49^. To test whether transcription is required to maintain SGO1 levels in oocytes, we used two mechanistically distinct RNA polymerase II inhibitors: triptolide, which blocks transcription initiation, and α-amanitin, which impairs elongation^60^. Both inhibitors effectively suppressed nascent transcription in early GV oocytes isolated from young mice, as confirmed by reduced incorporation of labeled nucleotides (Figure S1A). Additionally, we performed immunofluorescence to detect actively elongating RNA Pol II (RNA Pol II-pS2) and RNA FISH to detect major satellite (MajSat) repeat RNAs, which are expressed primarily from pericentromeric regions. Triptolide treatment resulted in a marked reduction of RNA Pol II-pS2 at centromeres and significantly decreased the levels of *MajSat* repeat RNAs at centromeres and pericentromeres in metaphase MI oocytes (Figures 2A–D).

**Figure 2.**
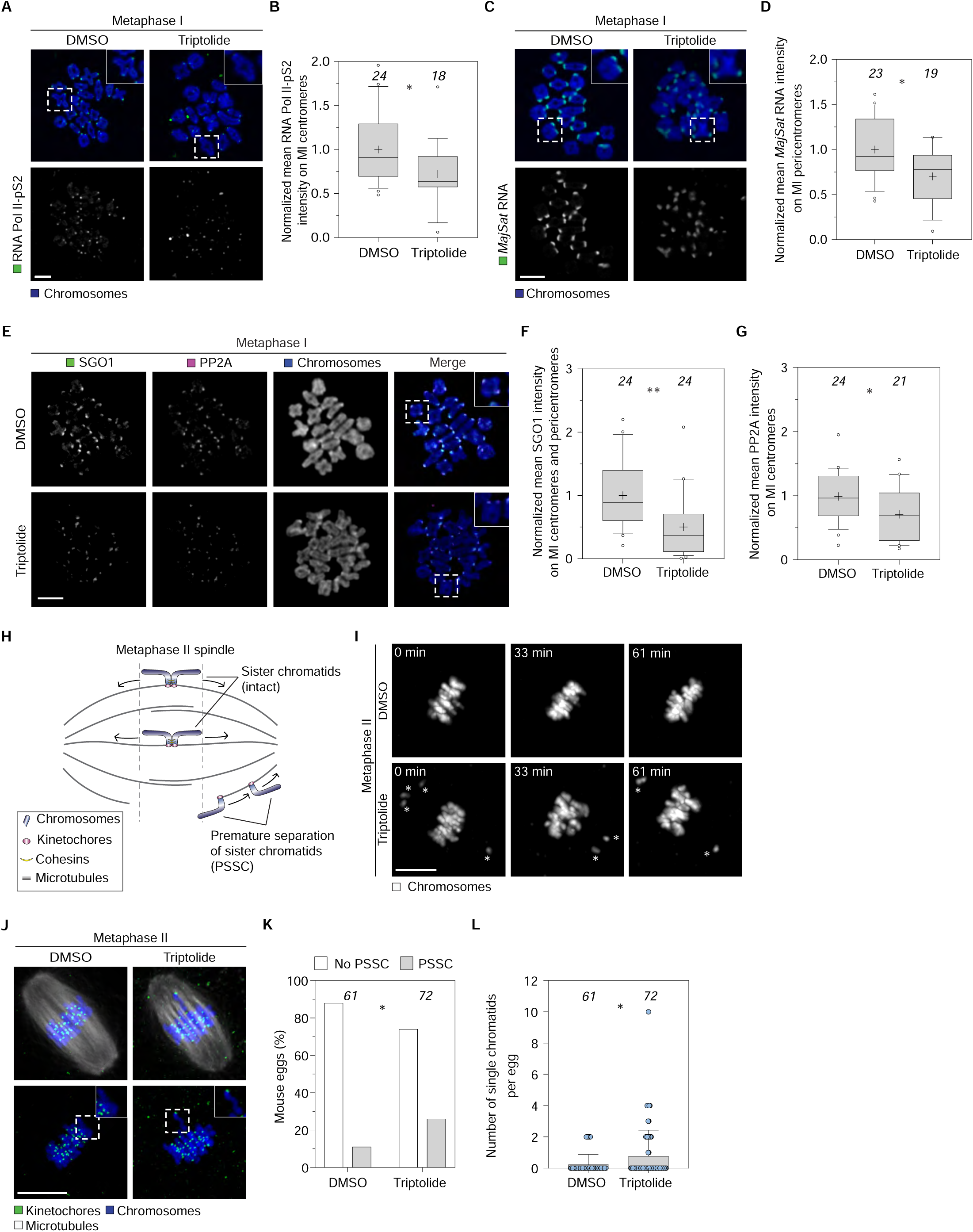
Pericentromeric transcription regulates SGO1/PP2A recruitment and prevents PSSC. (A–B) Pericentromeric transcription is reduced in triptolide-treated oocytes compared to controls. Representative images of metaphase I chromosomes (in blue) from oocytes treated with DMSO (left panel) and triptolide (right panel) with RNA Pol II-pS2 staining (green). White boxes with dashed lines indicate chromosomes at higher magnification in DMSO- and triptolide-treated oocytes. Relative intensity of the centromeric pool of actively elongating RNA Pol II from DMSO- and triptolide-treated oocytes, plots show n = number of oocytes analyzed at top, mean (plus), median (horizontal black line), 25th and 75th percentiles (boxes), 10th and 90th percentiles (whiskers), and the 1st and 99th percentiles (circles). statistical significance measured by Kolmogorov-Smirnov test, *p≤0.05. Scale bar: 10 µm. (C–D) Triptolide reduces *MajSat* RNA levels on metaphase I chromosomes. Representative RNA FISH images of metaphase I chromosomes (blue) from oocytes treated with DMSO (left panel) and triptolide (right panel) with *MajSat* RNA staining (green). The relative intensity of *MajSat* RNA at the pericentromere from DMSO- and triptolide-treated oocytes, plots shown, number of oocytes analyzed at top, statistical significance measured by Mann-Whitney test, *p≤0.05. Scale bar: 10 µm. (E–G) Representative immunofluorescence images of metaphase I chromosome spreads (blue) from control (DMSO-treated, top panel) and triptolide-treated (bottom panel) oocytes. (I–M) Triptolide reduces SGO1 (green) and PP2A (magenta) on metaphase I chromosomes compared to control. White boxes with dashed lines indicate chromosomes with higher magnification in DMSO- (upper right panel) and triptolide-treated (lower right panel) oocytes. Plots show n = number of oocytes analyzed at top, mean (plus), median (horizontal black line), 25th and 75th percentiles (boxes), 10th and 90th percentiles (whiskers), and the 1st and 99th percentiles (circles). statistical significance measured by Mann-Whitney test, *p≤0.05, **p≤0.01. Scale bar: 10 µm. (H) Schematic of sister chromatid alignment at the mouse metaphase II spindle. While sister chromatids are usually accurately aligned at the metaphase plate, prematurely separated chromatids are often misaligned due to unbalanced spindle microtubule tension. (I) Metaphase II chromosomes (grey) in DMSO (control) (upper panel) and triptolide-treated (lower panel) eggs imaged by time-lapse imaging at T= 0, 33, and 61 min at metaphase II. White asterisks indicate chromosomes with alignment defects, scored according to the scheme in (H). Scale bar: 10 µm. (J) Representative images of metaphase II chromosomes (blue) from eggs treated with DMSO (left panel) and triptolide (right panel). White boxes with dashed lines indicate chromosomes at higher magnification in DMSO and triptolide-treated eggs. Scale bar: 10 µm. (K) Frequency of PSSC measured by scoring the percentage of eggs with PSSC. Plots show number of eggs analyzed on top. Statistical significance is measured by Fisher’s exact test on the number of eggs with or without PSSC, *p≤0.05. (L) Number of single chromatids in eggs treated with DMSO and triptolide. Bars show the mean and standard deviation (black vertical line), individual values (blue dots). Statistical significance was measured by Kolmogorov-Smirnov test, *p≤0.05. See also Figure S1 and S2.

Notably, triptolide treatment significantly reduced SGO1 levels at centromeres and pericentromeres in metaphase I oocytes, while having no effect on SGO2 (Figures 2E, 2F, S2A, and S2B). PP2A levels were also reduced in metaphase I oocytes after triptolide treatment compared to controls (Figures 2E and 2G), consistent with loss of SGO1-mediated recruitment of PP2A. REC8 levels show a trend towards reduction in metaphase II but did not change during metaphase I, suggesting that cohesion maintenance prior to anaphase I is independent of transcription (Figures S2C–F). Furthermore, triptolide significantly reduced the intensity of the histone H3 variant CENP-A at centromeres in metaphase MII oocytes, as detected by immunofluorescence (Figures S2G and S2H). Centromeric and pericentromeric transcription has been implicated in loading CENP-A during G1 in mitotic cells^61–63^. Our data imply that CENP-A maintenance at centromeres in oocytes is transcription-dependent, given that triptolide treatment happened after presumed loading. We conclude that transcription is essential for maintaining high levels of SGO1 at the centromere and pericentromere, and of PP2A and CENP-A at the centromere.

Importantly, treatment with either triptolide or α-amanitin led to a substantial increase in PSSC, as well as increased alignment defects at the spindle equator in MII eggs compared to controls (Figures 2H, 2I, S1A, and S1B).

Overall, these data support a model in which active transcription and/or transcripts at pericentromeres are required to maintain SGO1-dependent and robust sister chromatid cohesion and centromere organization in oocytes from young mice.

### Triptolide-induced PSSC in young oocytes can be rescued with exogenous *MajSat* RNA

*MajSat* RNA colocalized with SGO1 at centromeric and pericentromeric regions, as determined by RNA FISH and immunofluorescence (Figure S1D). To determine if exogenous delivery of *MajSat* RNA could rescue the cohesion defects caused by transcriptional inhibition, we injected fluorescently labeled *MajSat* RNA into triptolide-treated GV oocytes isolated from young mice and monitored meiosis progression using live microscopy and immunofluorescence (Figure 3A). Exogenous *MajSat* RNA localized to centromeric and pericentromeric regions in live oocytes, while a control RNA did not (Figure 1C). Notably, *MajSat* RNA supplementation restored *MajSat* RNA (Figures 3B and 3C), SGO1 (Figures 3D and 3E), and PP2A (Figures 3D and 3F) levels on metaphase I chromosomes in triptolide-treated oocytes. Moreover, exogenous *MajSat* RNA significantly reduced the incidence of PSSC in triptolide-treated MII oocytes (Figures 3G and 3H). These results demonstrate that exogenous *MajSat* transcripts are sufficient to restore localized accumulation of SGO1 and PP2A and reduce PSSC in young oocytes subjected to transcriptional inhibition. Together, these data support a model in which pericentromeric RNA is required for Sgo1 function.

**Figure 3.**
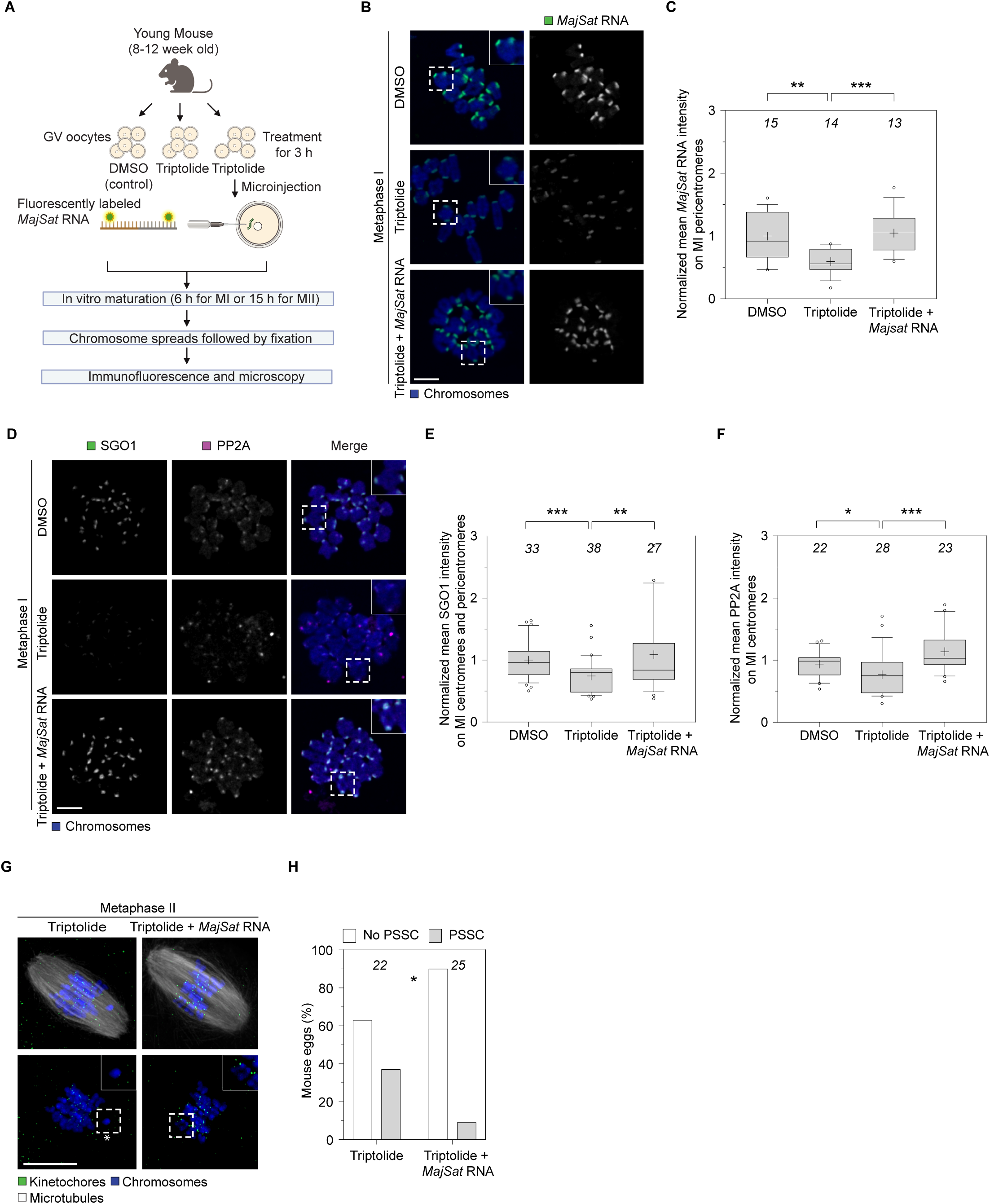
Decline in SGO1-PP2A recruitment upon triptolide treatment can be rescued by exogenous major satellite transcripts. (A) Schematic of chromosome error-reduction assay in mouse eggs upon triptolide treatment. Figure was created with BioRender.com. (B–C) Rescue of *MajSat* RNA levels in triptolide-treated oocytes after exogenous delivery of *MajSat* RNA. Representative images of metaphase I chromosomes (in blue) from DMSO-treated (top panel) and uninjected (middle panel) and *MajSat* RNA-injected (bottom panel) triptolide-treated oocytes with *MajSat* RNA staining (green). The relative intensity of *MajSat* RNA at the pericentromere from DMSO and both uninjected and injected triptolide-treated oocytes (B), plots shown, number of oocytes analyzed at top, statistical significance measured by Mann-Whitney test, **p≤0.01, ***p≤0.001. Scale bar: 10 µm. (D–F) Rescue of SGO1 and PP2A levels in triptolide-treated oocytes after exogenous delivery of *MajSat* RNA. Representative images of metaphase I chromosomes (in blue) from oocytes treated with DMSO (top panel) and uninjected (middle panel) and *MajSat* RNA injected (bottom panel) triptolide-treated oocytes. Chromosomes are immunostained with SGO1 (green) and PP2A (magenta). White boxes with dashed lines indicate chromosomes at higher magnification in oocytes in the insets. Plots show n = number of oocytes analyzed at top, mean (plus), median (horizontal black line), 25th and 75th percentiles (boxes), 10th and 90th percentiles (whiskers), and the 1st and 99th percentiles (circles). statistical significance measured by Mann-Whitney test. *p≤0.05, **p≤0.01, ***p≤0.001. Scale bar: 10 µm. (G) Representative images of metaphase II chromosomes (blue) of triptolide-treated eggs (left panel) and Triptolide-treated eggs with exogenous *Majsat*RNAs (right panel). White boxes with dashed lines indicate chromosomes at higher magnification in oocytes. Scale bar: 10 µm. (H) Frequency of PSSC is measured by scoring the number of single chromatids per egg. Plots show number of eggs analyzed on top. Statistical significance is measured by Fisher’s exact test on the number of eggs with or without PSSC, *p≤0.05.

### PSSC in aged oocytes is accompanied by a reduction of pericentromeric transcription, Sgo1, and PP2A

We observed higher PSSC frequencies and an increased number of single chromatids per egg in MII eggs from aged (60-65 weeks) versus young (8-12 weeks) mice (Figures 4A–C), consistent with a role for PSSC in age-dependent aneuploidy. The distance between sister kinetochores was also significantly increased in MII eggs from aged mice (Figure 4D).

**Figure 4.**
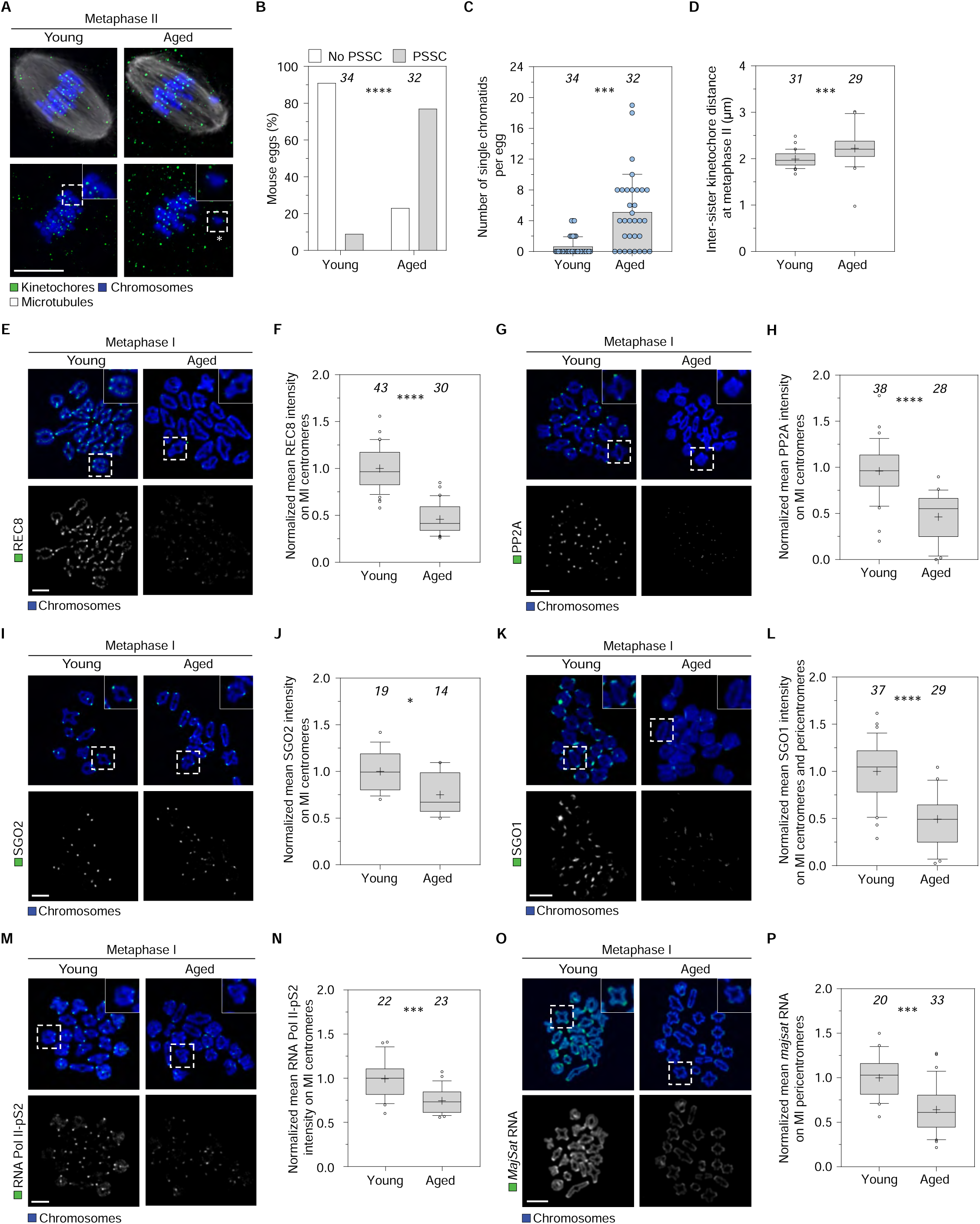
Age-related loss of SGO1, PP2A, and transcription drives sister chromatid cohesion failure. (A) Chromosomes (in blue) are properly aligned at the metaphase plate of young metaphase II eggs (top panel), but are prematurely separated in aged eggs (bottom panel). White boxes with dashed outlines show intact sister chromatids (upper inset panel) and separated sister chromatids (lower inset panel) in young and aged eggs at higher magnification. White asterisks indicate the presence of single chromatids. Scale bar: 10 µm. (B) The percentage of eggs with PSSCs in eggs from young (8-12 weeks old) and aged mice (60-65 weeks old). n, number of eggs analysed at top, statistical significance measured by Fisher’s exact test on the number of eggs with or without PSSC, ****p≤0.0001. (C) Frequency of PSSC measured by scoring the number of single chromatids per egg. Bars show the mean and standard deviation (black vertical line), individual values (blue dots). Statistical significance was measured using the Kolmogorov-Smirnov test, ***p≤0.001. (D) Age-related increase in interkinetochore distance. Plots show mean (plus), median (horizontal black line), 10th - 90th percentiles. Statistical significance was measured using the Mann-Whitney test, ***p≤0.001. (E–F) Representative images of metaphase I chromosomes (in blue) from young (top panel) and aged (bottom panel) oocytes with REC8 (green) staining. The relative intensity of the centromeric pool of REC8 from young and aged oocytes, plots show n = number of oocytes analysed at top, statistical significance measured by Kolmogorov-Smirnov test, ****p≤0.0001. Scale bar: 10 µm. (G–H) Representative images of metaphase I chromosomes (in blue) from young (top panel) and aged (bottom panel) oocytes with PP2A (green) staining. The relative intensity of centromeric PP2A from young and aged oocytes, plots show n = number of oocytes analysed on top, statistical significance measured by Kolmogorov-Smirnov test, ****p≤0.0001. Scale bar: 10 µm. (I–J) Representative images of metaphase I chromosomes (blue) from young (left panel) and aged (right panel) oocytes with SGO2 (green) staining. The relative intensity of the centromeric SGO2 from young and aged oocytes, plots show n = number of oocytes analysed on top, statistical significance measured by Kolmogorov-Smirnov test, *p≤0.05. Scale bar: 10 µm. (K–L) Representative images of metaphase I chromosomes (in blue) from young (top panel) and aged (bottom panel) oocytes with SGO1 (green) staining. The relative intensity of centromeric and pericentromeric SGO1 from young and aged oocytes, plots show n = number of oocytes analysed on top, statistical significance measured by Kolmogorov-Smirnov test, ****p≤0.0001. Scale bar: 10 µm. (M–P) Pericentromeric transcription is reduced in aged oocytes (left panel) compared to young oocytes (right panel), with reduced Pol II-pS2 RNA at the inner centromere (M–N) and reduced *MajSat* RNA at the pericentromeres (O–P). White boxes with dashed lines indicate chromosomes with higher magnification (M, O). Plots show n = number of oocytes analyzed at top, mean (plus), median (horizontal black line), 25th and 75th percentiles (boxes), 10th and 90th percentiles (whiskers), and the 1st and 99th percentiles (circles). statistical significance measured by Mann-Whitney test, ***p≤0.001. Scale bars: 10 µm. See also Figure S3.

To determine whether the cohesion protection mechanisms described above deteriorate with age, we analyzed chromosome spreads from MI oocytes. The cohesin component REC8 was diminished at centromeres in aged versus young oocytes, in line with previous reports^20–22^ (Figures 4E and 4F). We also observed significantly reduced levels of chromosome-associated PP2A, SGO1, SGO2, RNA Pol II-pS2, and *MajSat* RNA in aged MI oocytes (Figures 4G-P)^9,55,64^. Western blotting revealed reduced global levels of SGO1 in aged GV oocytes, while RNA Pol II-pS2 showed only a minor decrease when normalized to that of the loading control DDB1 (Figures S3A–C). The reduced global SGO1 abundance, accompanied by only minor changes in RNA Pol II-pS2, suggests that age-related centromere defects and PSSC arise primarily from the selective loss of factors that protect cohesion.

### PSSC in aged mouse eggs can be rescued by *Sgo1* supplementation

Having established that the levels of key cohesion-protective factors on chromosomes decline with age, we next examined whether reintroducing these factors in aged oocytes could reduce PSSC. We microinjected *MajSat* RNA, or mRNAs encoding SGO1 or SGO2, individually and in combination, into aged GV oocytes, followed by in vitro maturation (Figure 5A). Expression of *Sgo1* in aged oocytes was sufficient to reduce PSSC and restore inter-kinetochore distances at metaphase II, comparable to levels observed in young oocytes (Figures 5B–E). Combined administration of *Sgo1* with either *Sgo2* or *MajSat* RNA had similar effects (Figures S4A–D). In contrast, supplementation with either *Sgo2* or *MajSat* RNA alone had a smaller non-significant effect (Figures 5B–E and S4A–D). *MajSat* RNA alone is thus not sufficient to restore sister chromatid cohesion, likely because of limiting levels of SGO1 in aged oocytes.

**Figure 5.**
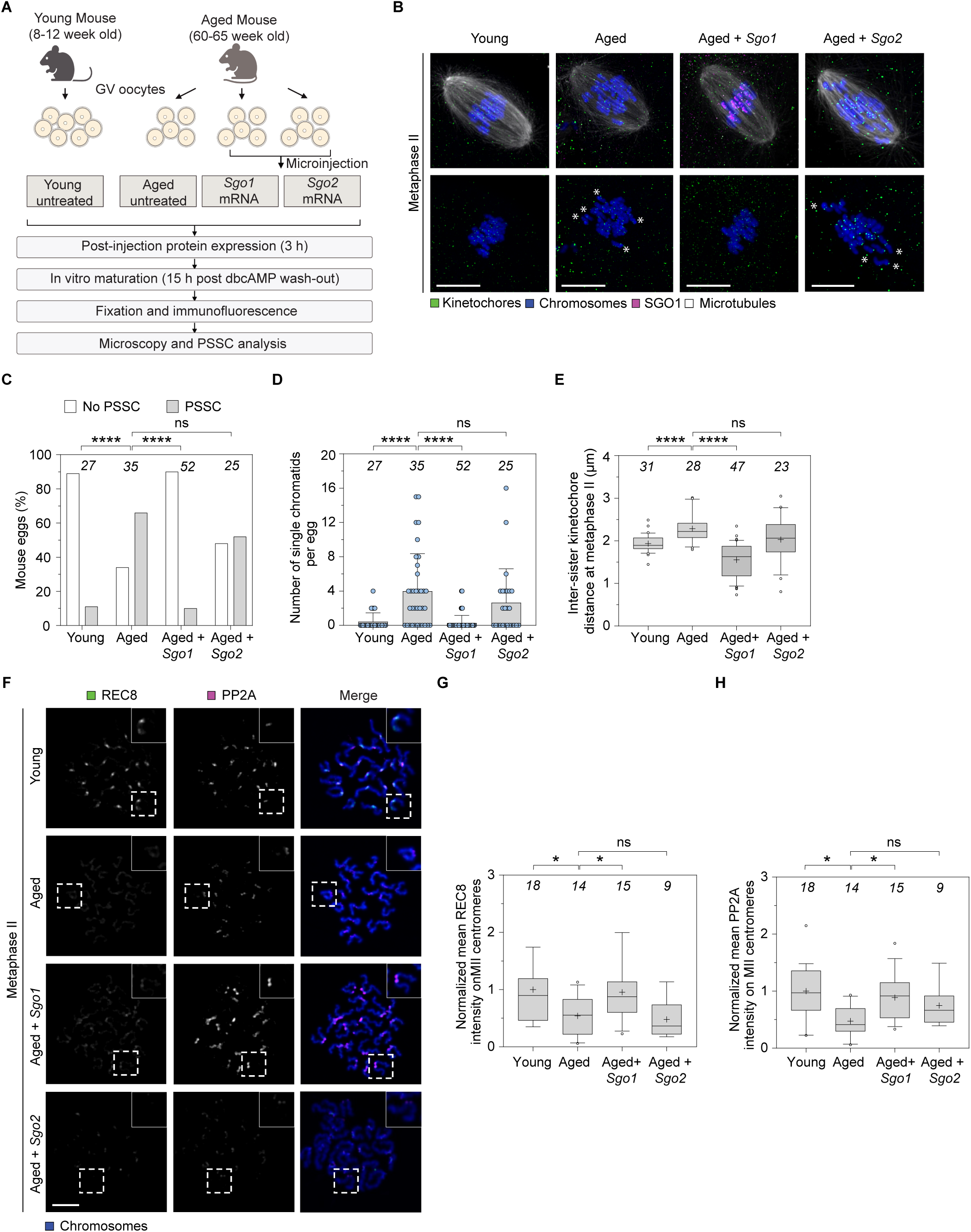
*Sgo1* supplementation rescues cohesion in aged mouse eggs. (A) Schematic of the assay for PSSC reduction in aged mouse eggs. Figure was created with BioRender.com. (B) *Sgo1* mRNA expression reduces PSSC in aged eggs compared to controls. Representative images of metaphase II chromosomes (in blue) from young and aged (untreated) eggs, aged eggs injected with *Sgo1* or *Sgo2* mRNA. White asterisks indicate the presence of single chromatids (lower panel). Scale bars: 10 µm. (C–D) Percentage of eggs with PSSC (C) and frequency of PSSC (D) in aged eggs supplemented with *Sgo1* or *Sgo2* mRNA. Plots show n = number of eggs analyzed at top, mean and standard deviation (black vertical line), individual values (blue dots). Statistical significance is measured by Fisher’s exact test on the number of eggs with or without PSSC (C) and Kolmogorov-Smirnov test (D). ns = not significant, ****p≤0.0001. (E) Measurements of interkinetochore distances between sister chromatids of metaphase II chromosomes in mouse eggs injected with the combination of RNAs indicated in (B). Plots show n = number of eggs analyzed at top, mean (plus), median (horizontal black line), 25th and 75th percentiles (boxes), 10th and 90th percentiles (whiskers), and the 1st and 99th percentiles (circles). Statistical significance was measured by Kolmogorov-Smirnov test. ns = not significant, ****p≤0.0001. (F) Representative images of metaphase II chromosomes (in blue) immunostained with REC8 (in green) and PP2A (in magenta) from young and aged untreated eggs and from aged eggs supplemented with *Sgo1* or *Sgo2* mRNA. Scale bar: 10 µm. (G–H) *Sgo1* mRNA rescues centromeric PP2A and REC8 levels in aged metaphase II eggs, but *Sgo2* does not. White dashed boxes show chromosomes at higher magnification. Plots show n = number of oocytes analyzed at top, mean (plus), median (horizontal black line), 25th and 75th percentiles (boxes), 10th and 90th percentiles (whiskers), and the 1st and 99th percentiles (circles). Statistical significance is measured by Kolmogorov-Smirnov test. ns = not significant, *p≤0.05. See also Figure S4.

To determine if reintroducing *Sgo1* also increases REC8 and PP2A at centromeres, we performed immunofluorescence on chromosome spreads. Aged GV oocytes supplemented with *Sgo1* showed increased levels of REC8 and PP2A on metaphase chromosomes at MI and MII (Figures 5F–H and S4E–G). In contrast, *Sgo2* supplementation in aged GV oocytes did not lead to a significant increase in chromosomal PP2A levels at MII (Figures 5F–H and S4E–G). Notably, supplementation with the *Sgo1* N61I mutant, which is unable to bind PP2A, failed to rescue PSSC (Figures 4A–D). Overall, these data indicate that the reintroduction of *Sgo1* in aged oocytes can prevent PSSC through recruitment of PP2A^31,65^.

### *SGO1* supplementation reduces PSSC in human eggs

Human oocytes exhibit a high incidence of PSSC with age, a principal driver of aneuploidy and IVF failure. To determine whether SGO1 is associated with an age-dependent increase in PSSC in human oocytes, we examined SGO1 localization and function in eggs donated from women of different ages. SGO1 localized to an extended region spanning centromere and pericentromere at metaphase I in human oocytes (Figure 6A), consistent with our observations in mouse oocytes (Figure 4K) and previous reports^21,66^. Importantly, chromosomal levels of SGO1 were significantly reduced in oocytes from women aged ≥35 years compared to those from women aged <35 years (Figures 6A, 6B; Table S2). The data thus indicate an age-related decline in this key cohesion protector. Notably, the frequency of PSSC was significantly higher among chromosomes and metaphase II eggs from women aged 35 years and older compared to younger women (Figures 6C and 6D). We infer that loss of SGO1, accompanied by an increase in PSSC, is a conserved hallmark of oocyte aging in both mice and humans.

**Figure 6.**
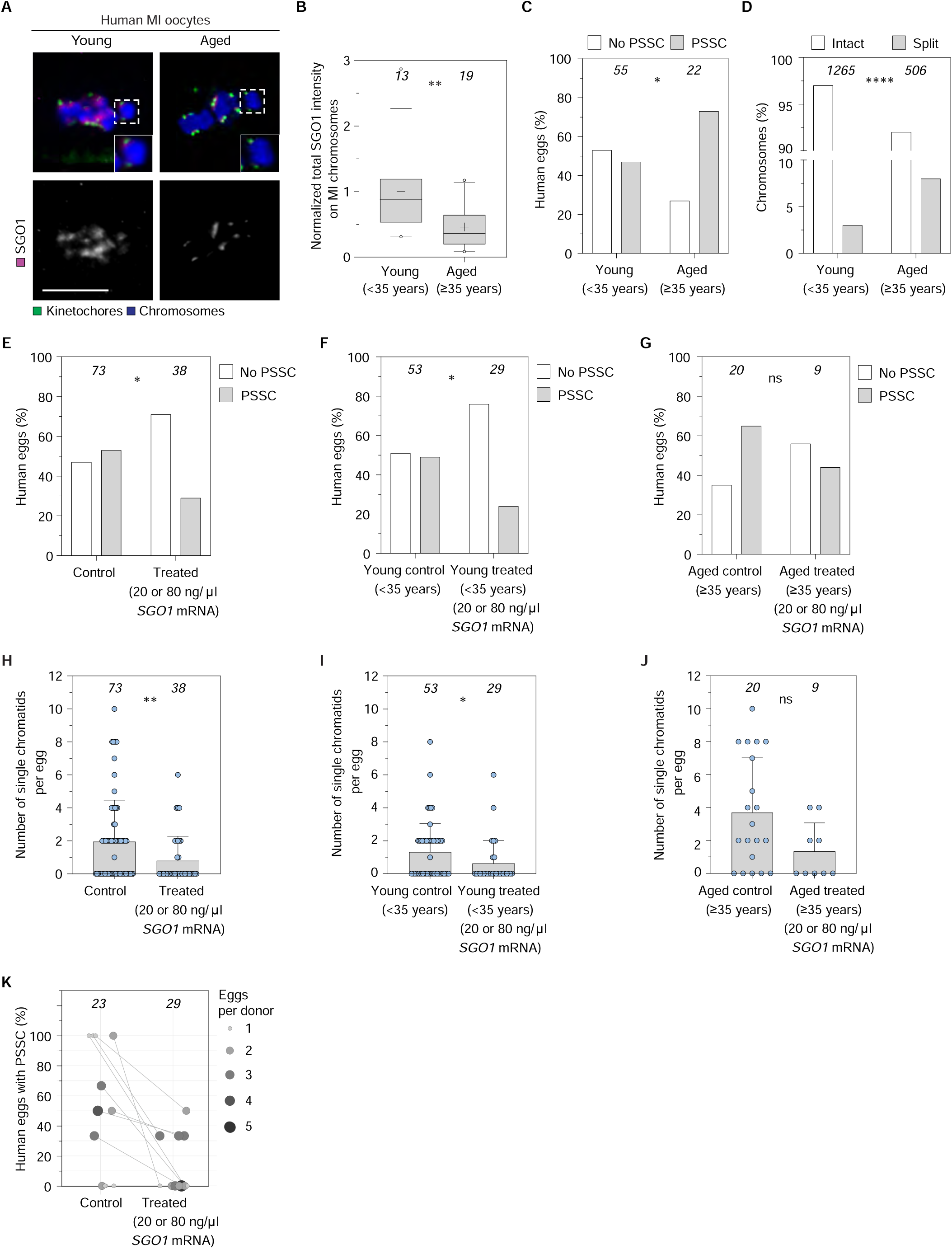
SGO1 levels decline in aged human oocytes and supplementation rescues PSSC. (A) Representative images showing loss of SGO1 at the centromere and pericentromere region of human metaphase I oocytes from aged (≥35 years) women compared to young (<35 years). White dashed boxes show chromosomes at higher magnification. SGO1 (magenta), kinetochores (CREST, green), and chromosomes (Hoechst, blue) are shown. Scale bar: 10 µm. (B) Normalized total intensity of chromosomal SGO1 in metaphase I oocytes from young (<35 years) and aged (≥35 years) women. SGO1 is significantly reduced at the pericentromere region in aged human oocytes compared to young oocytes. Plots show n = number of oocytes analysed at top, mean (plus), median (horizontal black line), 25th and 75th percentiles (boxes), 10th and 90th percentiles (whiskers), and the 1st and 99th percentiles (circles). Statistical significance is measured by the Mann-Whitney test, **p≤0.01. (C–D) Frequency of PSSC as measured by the percentage of eggs and chromosomes with PSSC is significantly increased in eggs from aged (≥35 years) women in comparison with those from young (<35 years) women. Plots show n = number of eggs (C) and chromosomes (D) analyzed at top. Statistical significance is measured by Fisher’s exact test on the number of eggs (C) and chromosomes (D) with or without PSSC. *p≤0.05, ****p≤0.0001. Scale bars: 10 µm. (E–G) Eggs from young (<35 years) and aged (≥35 years) women were microinjected with 4 pl of *SGO1* mRNA (20 or 80 ng/µl) or kept untreated. Treated and untreated eggs were scored for PSSC. The percentage of eggs with PSSC is shown for young and aged women together (E), as well as for the same young (<35 years, F) and aged (≥35 years, G) women, separately. Plots show n = number of eggs analyzed at top. Statistical significance is measured by Fisher’s exact test on the number of eggs with or without PSSC. *p≤0.05, ns = not significant. (H–J) *SGO1* mRNA expression (20 or 80 ng/µl) reduces frequency of PSSC in human eggs. Frequency of PSSC is measured by scoring the number of single chromatids per egg in untreated eggs and eggs supplemented with 4 pl of 20 or 80 ng/µl *SGO1* mRNA (young and aged eggs together (h), in eggs from young (<35 years, i) and aged (≥35 years, j) women. Bars show the mean and standard deviation (black vertical line), individual values (blue dots). Plots show n = number of chromosomes analyzed at top. Statistical significance is measured by Mann-Whitney test, ns = not significant, *p≤0.05, **p≤0.01. (K) Eggs from the same donor were microinjected with 4 pl of *SGO1* mRNA (20 or 80 ng/µl) or kept untreated. The percentage of PSSC in the treated and untreated eggs is shown for each donor (linked by a line). The size of the circle corresponds to the number of eggs that was analyzed. Plots show n = number of eggs analyzed at top, each dot represents percentage of eggs with PSSC per donor. See also Figure S5, S6, and table S2.

We hypothesized that *SGO1* supplementation could improve sister chromatid cohesion in human eggs. We first verified that *SGO1-EGFP* mRNA microinjected into donated human GV and MI oocytes led to the accumulation of SGO1-EGFP to the region spanning centromere and pericentromere at metaphase in both MI and MII human oocytes (Figure S5A). Next, we injected oocytes with two different doses of *SGO1* mRNA (4 pl of 20 or 80 ng/μl). The combined analysis across all donated eggs, exogenous SGO1 significantly reduced the proportion of eggs exhibiting PSSC, from 53% in controls to 29% in treated oocytes (Figure 6E). Supplementation of human eggs with exogenous *SGO1* thus nearly halved the fraction of eggs with PSSC. When stratified by age, the effect was statistically significant in eggs from women <35 years, in which the fraction of PSSC-positive eggs decreased from ∼49% to ∼24% following Sgo1 expression (Figure 6F). In eggs from women ≥35 years, a similar directional trend was observed (∼65% to ∼44%) (Figure 6G). However, this did not reach statistical significance (Figure 6G), likely reflecting the small number of treated oocytes (n = 9). Nevertheless, the possibility of an age-dependent difference in responsiveness cannot be excluded.

We next assessed chromosome-level cohesion. Across all donors, *SGO1* supplementation significantly reduced the number of single chromatids per egg (Figure 6H). When stratified by age, oocytes from younger donors should a significant reduction of single chromatids per egg following *SGO1*-treatment (Figure 6I). Oocytes from women ≥35 years exhibited a markedly higher frequency of single chromatids, and *SGO1* expression produced a similar directional trend toward fewer single chromatids, although this did not reach statistical significance (Figure 6J). Together, these data indicate that *SGO1* supplementation improves centromeric cohesion in human eggs and can partially restore chromosome-level cohesion in eggs from older women.

To rigorously assess the therapeutic potential of *SGO1* supplementation, we compared untreated eggs with *SGO1*-treated eggs from the same donors. In every paired comparison, *SGO1* supplementation reduced the percentage of eggs with PSSC (Figure 6K). These within-donor analyses demonstrate that SGO1 supplementation consistently increases the fraction of PSSC-free eggs at the level of individual patients.

Combined analysis of both *SGO1* mRNA doses revealed a consistent reduction in PSSC and single - chromatid defects. Dose-stratified analyses showed that supplementation of oocytes with *SGO1* at both 20 and 80 ng/µL resulted in significantly increased chromosomal SGO1 Levels at metaphase II (Figure S5B). When analyzed by dose, rescue of PSSC in metaphase II eggs and individual chromosomes was most pronounced at 20 ng/µL (Figure 5C–J).

Together, our findings demonstrate that age-dependent loss of SGO1 contributes to premature sister chromatid separation and chromosome segregation errors in mammalian oocytes, and that restoring SGO1 levels can preserve centromeric cohesion and reduce these errors in both mouse and human eggs (Figure S6A). These results establish a previously unrecognized mechanism of cohesion deprotection during oocyte aging and identify *SGO1* supplementation as a potential means to improve chromosome segregation fidelity in human eggs.

## Discussion

Aneuploidy arising from errors in chromosome segregation is the principal cause of age-related infertility, IVF failure, and miscarriage in women, yet no therapeutic interventions have been available to address this problem in humans^1,6,7^. Although several experimental approaches have been shown to reduce premature sister chromatid separation in mouse oocytes, whether cohesion defects could be corrected in human eggs has remained unknown^53,54^.

Our study identifies loss of the cohesin protector Shugoshin 1 (SGO1) as a key contributor to premature sister chromatid separation (PSSC) and, therefore, aneuploidy in aged mammalian eggs. Moreover, we demonstrate that *Sgo1* supplementation can restore chromatid cohesion and reduce PSSC in both mouse and human eggs. This work not only elucidates a previously unrecognized mechanism underlying age-related cohesion loss but also provides the first evidence that targeted molecular intervention can improve chromosomal integrity in aged human eggs. Our findings indicate that loss of SGO1 in aged oocytes contributes to the reduction of centromeric and pericentromeric cohesion and to the increased incidence of premature sister chromatid separation.

Although SGO2 is known to be essential for sister chromatid cohesion in young oocytes^28,58^ and is depleted from pericentromeric regions in aged human eggs^55^, supplementation of *Sgo2* was insufficient to restore cohesion or reduce PSSC in aged mouse oocytes. In contrast, *Sgo1* supplementation robustly restored cohesion, reducing PSSC to levels observed in young oocytes. During meiotic maturation, SGO1 localizes to a broader domain spanning the centromere and pericentromere than SGO2, consistent with a distinct role in stabilizing sister chromatid cohesion. Moreover, the SGO1 N61I mutant, which abolishes PP2A binding, fails to rescue cohesion, indicating that SGO1 acts through PP2A recruitment to protect cohesin.

A key advance of this study is the demonstration that transcription is essential for maintaining SGO1 and PP2A at meiotic centromeres. Transcription inhibition leads to the loss of both factors, and increased PSSC. In young oocytes, exogenous major satellite RNA can localize to pericentromeric regions and partially restore the defects caused by transcriptional inhibition. However, supplementation with satellite RNA alone is insufficient to rescue cohesion in aged oocytes, indicating that the age-related loss of SGO1 protein is a limiting factor that cannot be overcome by restoring transcripts alone. These findings highlight a critical interplay between transcriptional regulation and protein stability in the maintenance of chromosomal cohesion.

Importantly, we extend these mechanistic insights to human oocytes, where we find that SGO1 levels are significantly reduced in eggs from women of advanced maternal age. *SGO1* supplementation in human eggs consistently reduced PSSC and restored chromatid cohesion. The rescue effect was robust across donors, underscoring the translational potential of *SGO1*-based interventions. This represents the first demonstration that a molecular intervention can directly correct a defect in human egg chromosome cohesion, addressing a major unmet need in reproductive medicine.

In summary, our study reveals new mechanisms underlying age-related aneuploidy in mammalian eggs, providing a foundation for the development of targeted therapeutic strategies to improve egg quality for women undergoing assisted reproduction. By demonstrating that *SGO1* supplementation can restore chromatid cohesion in both mouse and human eggs, we open new avenues for fertility preservation and the prevention of maternal age-related chromosomal disorders.

## Acknowledgements

We are grateful to the staff of the Animal Facility and the Live-Cell Imaging Facility of the Max Planck Institute for Multidisciplinary Sciences for technical assistance. We thank the patients who participated in this study; the clinicians, and embryologists at the Reprofit International, Clinic for Reproductive Medicine, Brno, Czech Republic, and Bourn Hall Fertility Clinic; Bourn, United Kingdom, for their support of this study; E. Mönnich and A. Politi for advice on image analysis; M. Daniel, K. Rentsch, and L. Timm for coordinating the animal experiments; C. Thomas for helpful discussions about the project; A. Webster for generating the rabbit anti-REC8 antibody; Y. Watanabe for the rabbit anti-SGOL1 antibody; N. Sharma for generating the *hSGO1* construct, M. Lidschreiber and J. A. Garrido for their advice and exploratory work related to SGO1 supplementation strategies in mammalian oocytes; T. Padilla for the *MajSat* construct and J. A. Burton for his advice on generating *MajSat* RNA. Biorender.com was used as indicated in figure legends.

## Author contributions

D.S. and M.S. conceived the study. D.S. and M.S. designed experiments and methods for data analysis. S.M. performed microinjection of human oocytes at Bourn Hall Clinic. D.S. performed all other experiments and analyzed the data. T.C. performed some of the preliminary experiments on oocyte transcription and contributed in the initial phase of the project. A.P.Z. performed experiments in human oocytes used in Figure 6B to analyse age-related decline in chromosomal SGO1 level. Z.H., B.M., M.B., and K.E. supervised the collection of human oocytes. L.W. coordinated the collaboration with the IVF clinics and managed ongoing regulatory compliance for human oocyte work. D.S. and M.S. drafted the manuscript and prepared the figures with input from all authors. M.S. supervised the study.

## Supplemental figure titles and legends

**Figure S1.**
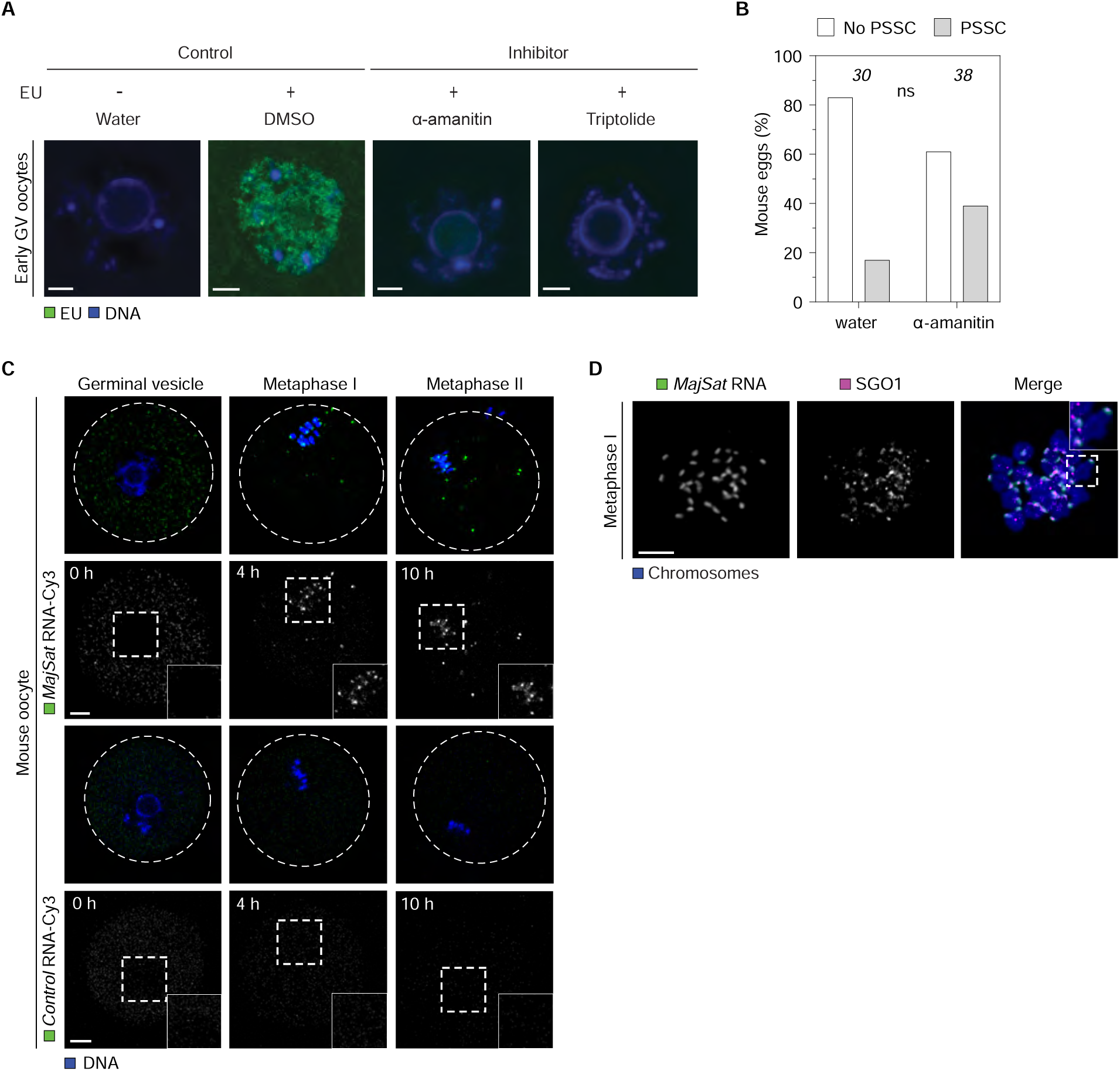
Active pericentromeric transcription during meiotic divisions in mouse oocytes. (A) Representative immunofluorescence images showing the localization nascent transcripts (EU, green) inside the nucleus (blue) of early GV oocytes. Images show EU levels in negative control (no EU), control (with EU), upon treatment with α-amanitin, and triptolide. Scale bar: 10 µm. (B) Frequency of PSSC with α-amanitin measured by scoring the percentage of eggs with PSSC, similar to levels observed with triptolide (Figure 2K). Plots show number of eggs analyzed on top. Statistical significance is measured by Fisher’s exact test, ns = not significant. (C) Representative time-lapse images showing the localization of *MajSat* RNA in GV and MI oocytes and MII eggs upon microinjection of GV oocytes with UTP-X-Cy3 labeled *MajSat* (top panel) and control (bottom panel) RNA. *MajSat* RNA localizes to the pericentromeres during metaphase I and metaphase II stages along with foci present in the cytoplasm. Cy3-labeled control RNA localizes only in the cytoplasm and not on chromosomes. *MajSat* and control RNA (green), chromosomes (H2B-SNAP, blue) are shown. Scale bars: 10 µm. (D) Co-localization of *MajSat* RNA and SGO1 observed on metaphase I chromosome spreads upon performing RNA FISH with immunofluorescence. Scale bar: 10 µm.

**Figure S2.**
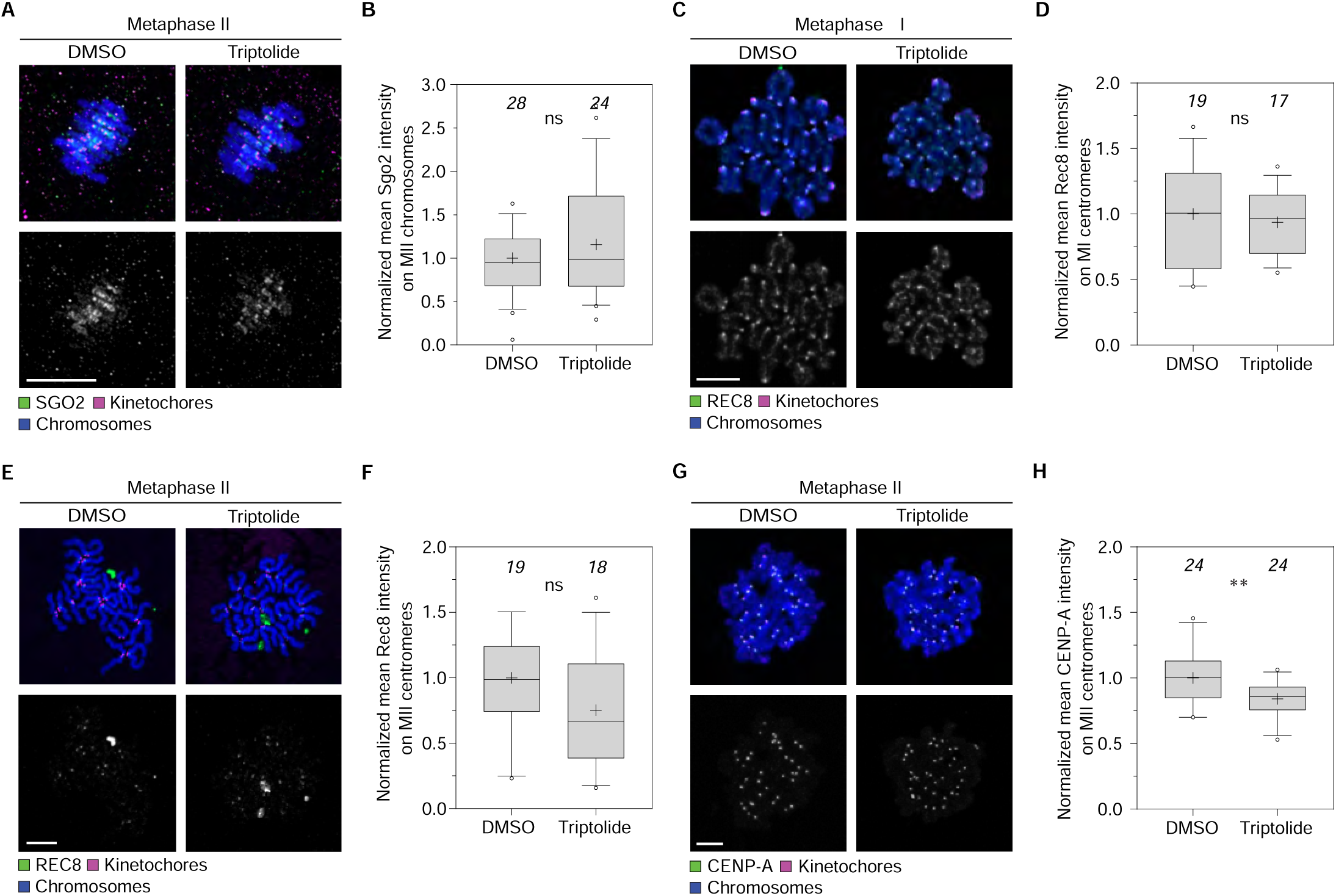
Triptolide reduces CENP-A levels at mouse centromeres, but not SGO2. (A–B) Representative images of metaphase II chromosomes (blue) immunostained for SGO2 (green) and kinetochores (CREST, magenta) from DMSO- and triptolide-treated eggs. Plots shown n = number of eggs analyzed at top, mean (plus), median (horizontal black line), 25th and 75th percentiles (boxes), 10th and 90th percentiles (whiskers), and the 1st and 99th percentiles (circles). Statistical significance is measured by Mann-Whitney test, ns = not significant. Scale bar: 10 µm. (C–F) Representative images of metaphase I (C) and metaphase II (E) chromosomes (blue) immunostained for REC8 (green) and kinetochores (CREST, magenta) from DMSO- and triptolide-treated metaphase I oocytes (C) and metaphase II eggs (E). Plots shown n = number of oocytes analyzed at top, mean (plus), median (horizontal black line), 25th and 75th percentiles (boxes), 10th and 90th percentiles (whiskers), and the 1st and 99th percentiles (circles). Statistical significance is measured by Mann-Whitney test, ns = not significant. Scale bars: 10 µm. (G–H) Triptolide reduces CENP-A at metaphase II centromeres. Plots show n = number of eggs analyzed at top, mean (plus), median (horizontal black line), 25th and 75th percentiles (boxes), 10th and 90th percentiles (whiskers), and the 1st and 99th percentiles (circles). Statistical significance is measured by Mann-Whitney test, **p≤0.01. Scale bar: 10 µm.

**Figure S3.**
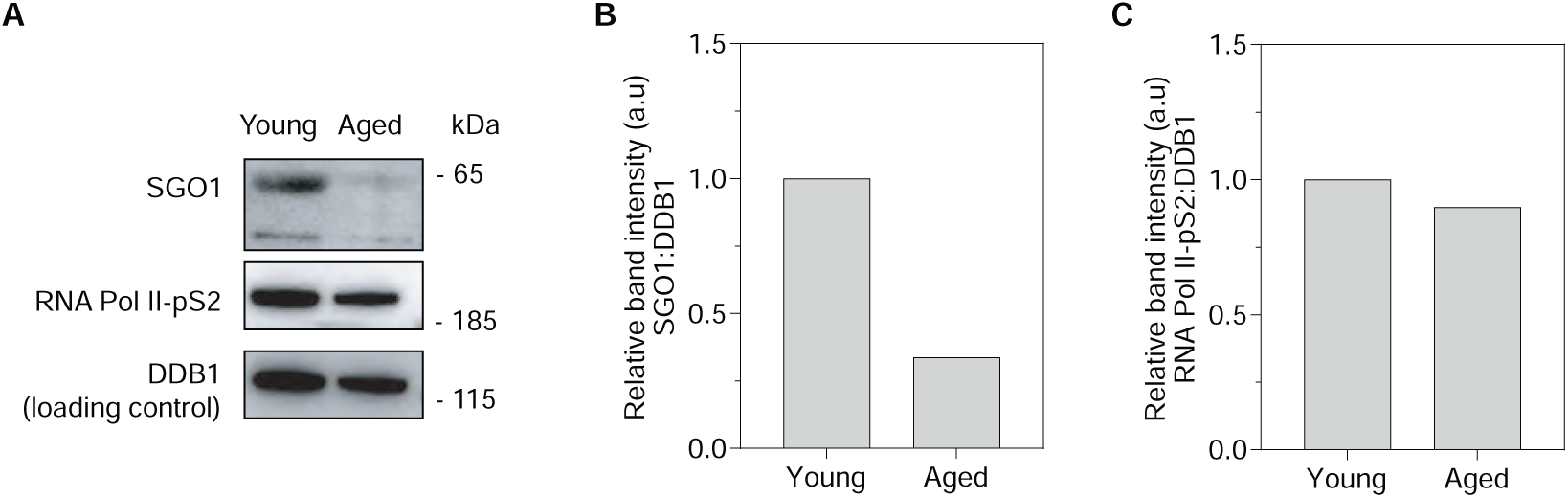
SGO1 protein levels reduce in aged mouse GV oocytes. (A) Levels of SGO1, RNA Pol II-pS2, and DDB1 observed on immunoblots of young and aged oocytes at the germinal vesicle stage. SGO1 protein levels show a pronounced reduction in aged oocytes, whereas RNA Pol II-pS2 shows a partial reduction compared to the loading control (DDB1). (B–C) Relative band intensity of SGO1 and RNA Pol II-pS2 in young versus aged oocytes upon normalization with Ddb1. SGO1 protein levels show a pronounced reduction in aged oocytes, whereas RNA Pol II-pS2 shows only a minor reduction.

**Figure S4.**
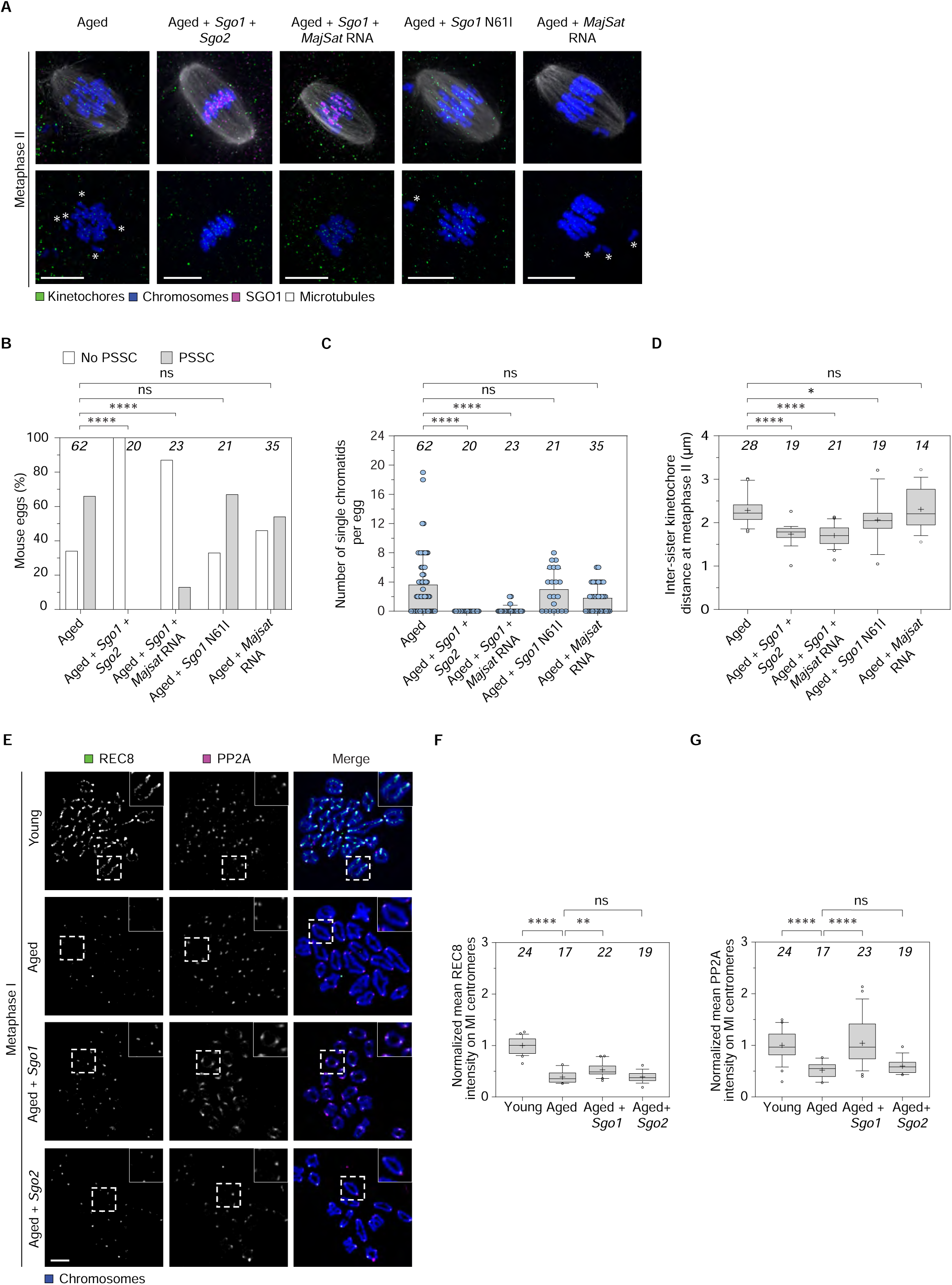
Rescue of cohesion and REC8/PP2A levels in aged mouse oocytes. (A) Representative images of metaphase II chromosomes (in blue) from aged (untreated) eggs (left panel), aged eggs supplemented with *Sgo1* together with *Sgo2*, *Sgo1*together with *MajSat* RNA, *Sgo1* N61I mutant, and *MajSat* RNA alone. Scale bars: 10 µm. (B–C) Percentage of eggs with PSSC (B) and frequency of PSSC (C) in aged eggs supplemented with *Sgo1* together with *Sgo2*, *Sgo1* together with *MajSat* RNA, *Sgo1* N61I mutant, and *MajSat* RNA alone. Scatter plots show n = number of eggs analyzed at top, mean and standard deviation (black vertical line), individual values (blue dots). Statistical significance is measured by Fisher’s exact test on the number of eggs with or without PSSC (B) and Kolmogorov-Smirnov test (C). ns = not significant, ****p≤0.0001. (D) Measurements of interkinetochore distances between sister chromatids of metaphase II chromosomes in mouse eggs injected with the combination of RNAs indicated in (A). Plots show n = number of eggs analyzed at top, mean (plus), median (horizontal black line), 25th and 75th percentiles (boxes), 10th and 90th percentiles (whiskers), and the 1st and 99th percentiles (circles). Statistical significance was measured by Kolmogorov-Smirnov test. ns = not significant, *p≤0.05, ****p≤0.0001. (E) Representative images of metaphase I chromosomes (blue) immunostained for REC8 (green) and PP2A (magenta) from young and aged untreated oocytes and aged oocytes supplemented with *Sgo1* or *Sgo2*. Scale bar: 10 µm. (F–G) *Sgo1* rescues centromeric REC8 and PP2A levels in aged metaphase I oocytes, but *Sgo2* does not. White dashed boxes show chromosomes at higher magnification. Plots show n = number of oocytes analyzed at top, mean (plus), median (horizontal black line), 25th and 75th percentiles (boxes), 10th and 90th percentiles (whiskers), and the 1st and 99th percentiles (circles). Statistical significance is measured by Kolmogorov-Smirnov test. ns = not significant, **p≤0.01, ****p≤0.0001.

**Figure S5.**
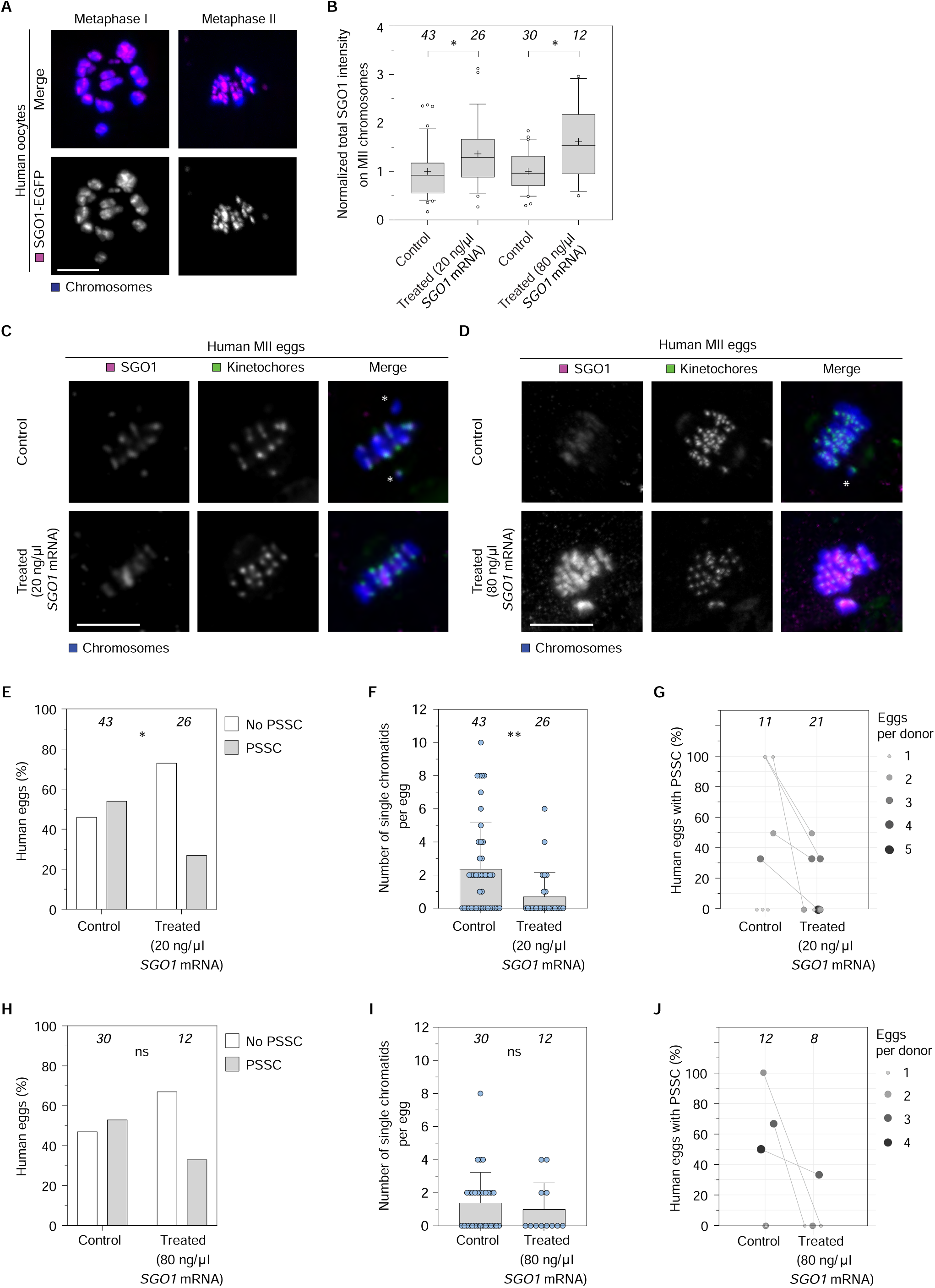
SGO1-mediated cohesion rescue is conserved across human donors and *SGO1* mRNA concentrations. (A) Representative images showing exogenous hSGO1-EGFP (magenta) localizes to human inner centromere and pericentromere bridge at metaphase I (with background signal on chromosome arms), and metaphase II. Scale bar: 10 µm. (B) Normalized total intensity of chromosomal SGO1 in metaphase II eggs microinjected with 4 pl of 20 (Treated 80 ng/μl) or 80 ng/µl SGO1 mRNA (Treated 80 ng/μl) or kept untreated (Control). Chromosomal SGO1 is significantly increased in eggs treated with 4 pl of 20 or 80 ng/µl SGO1 mRNA compared to controls. Plots show n = number of oocytes analysed at top, mean (plus), median (horizontal black line), 25th and 75th percentiles (boxes), 10th and 90th percentiles (whiskers), and the 1st and 99th percentiles (circles). Statistical significance is measured by the Mann-Whitney test, ns = not significant, *p≤0.05. (C–D) Immunofluorescence confocal images of human metaphase II eggs labelled for chromosomes (blue; Hoechst) and immunostained for SGO1 (magenta), and kinetochores (green; CREST) from untreated eggs (control) and treated eggs microinjected with either 4 pl of 20 ng/µl (Treated 20 ng/µl, c) or 80 ng/µl *SGO1* mRNA (Treated 80 ng/µl, d). SGO1 expression reduces PSSC in human eggs. White asterisks indicate the presence of single chromatids in control oocytes (upper panel). Scale bars: 10 µm. (E–F) Eggs were microinjected with 4 pl of *SGO1* mRNA at concentrations of 20 ng/µl, or kept untreated. Treated and untreated eggs were scored for PSSC. The percentage of eggs with PSSC (E) and number of single chromatids per egg (F) is shown. Plots show n = number of eggs analyzed at top. Statistical significance is measured by Fisher’s exact test on the number of eggs with or without PSSC (E) and Mann-Whitney test (F). *p≤0.05, **p≤0.01. (H–I) Eggs were microinjected with 4 pl of SGO1 mRNA at 80 ng/µl, or kept untreated. Treated and untreated eggs were scored for PSSC. The percentage of eggs with PSSC (H) and number of single chromatids per egg (I) is shown. Plots show n = number of eggs analyzed at top. Statistical significance is measured by Fisher’s exact test on the number of eggs with or without PSSC (H) and Mann-Whitney test (I). ns = not significant. (G, J) Eggs from the same donor were microinjected with 4 pl of 20 ng/µl (G) or 80 ng/µl (J) *SGO1* mRNA. The percentage of PSSC in the treated and untreated eggs is shown for each donor (linked by a line). The size of the circle corresponds to the number of eggs that was analyzed. Plots show n = number of eggs analyzed at top, each dot represents percentage of eggs with PSSC per donor. Scale bars: 10 µm.

**Figure S6.**
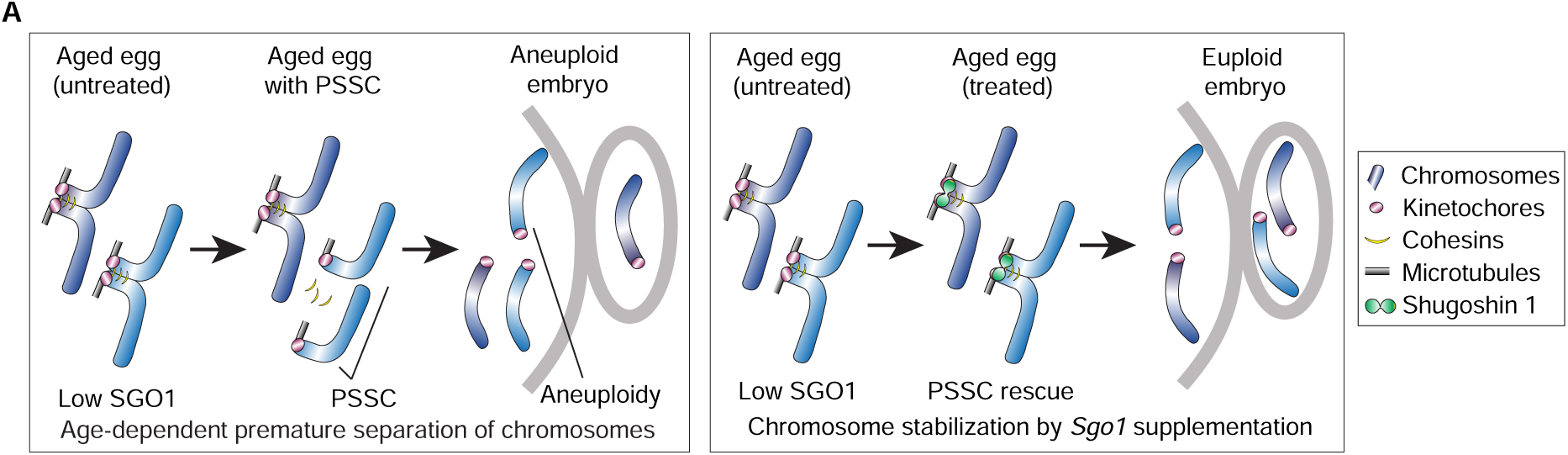
*Sgo1* rescues PSSC in aged eggs. (A) Schematic representation of error-prone chromosome segregation during meiotic progression in aged oocytes leading to PSSC and aneuploidy. *Sgo1* expression can rescue PSSC errors in aged eggs by protecting sister chromatid cohesion, thereby reducing aneuploidy.

## Methods

### Mouse oocyte and follicle isolation and culture

All mice were maintained in a specific pathogen-free environment and handled according to the guidelines of the Max Planck Institute for Multidisciplinary Sciences Animal House in accordance with international animal welfare standards (guidelines and recommendations of the Federation of Laboratory Animal Science Associations) and the German Animal Welfare Act (TSchG). Young (8-12-week-old) and aged (60-65-week-old) 129S6 mice were sacrificed by cervical dislocation and ovaries were collected in M2 medium supplemented with 750 μM dibutyryl cyclic AMP (dbcAMP, Sigma-Aldrich, #D0627) to maintain oocytes in prophase I arrest. Isolation and selection of GV stage oocytes (around 70 µm in diameter) with a central nucleus were performed after puncturing mouse ovaries with a hypodermic needle. GV oocytes were cultured in M2 media supplemented with dbcAMP under paraffin oil (NidaCon, #NO-400K) at 37°C. For meiotic maturation experiments, oocytes were washed out of dbcAMP.

For follicle isolation, 10-12-day old B6CBAF1 females were used as previously described with some modifications^59^. Ovaries were collected in MEM-alpha HEPES GlutaMAX (ThermoFisher, #42360024) supplemented with 0.1x penicillin-G/streptomycin (Gibco). Ovaries were briefly dissociated with 2 mg/ml Collagenase IV (Thermo Fisher Scientific, #17104019) for 5 min and then mechanically pipetted several times. Follicles were washed free of collagenase and selected to be approximately 100 µm in size and surrounded by uniform layers of granulosa cells. Follicle-enclosed oocytes were microinjected and plated on collagen-coated inserts (Corning, #3491/3493) with MEM-alpha GlutaMAX (ThermoFisher, #41090028) supplemented with 5% fetal bovine serum (FBS) (Gibco, #16000044), 1x insulin-transferrin-selenium (ITS, ThermoFisher Scientific, #41400045), 0.01 µg/ml ovine follicle stimulating hormone, oFSH (National Hormone and Peptide Program, #NDDK-oFSH-20) and 0.1x penicillin-G/streptomycin (Gibco, #15070-063) at 37°C/5% CO2. The medium in the wells surrounding the inserts was changed every 3 days. After 9-10 days of in vitro culture, GV oocytes were harvested and cultured in modified M2 medium containing 10% FBS instead of 4 mg/ml bovine serum albumin (BSA) until metaphase II stage.

### Preparation and culture of fresh and vitrified human oocytes

All human oocytes used in this study were obtained from Reprofit International Clinic for Reproductive Medicine (Brno, Czech Republic), and Bourn Hall Fertility Clinic (Bourn, United Kingdom). The use of unfertilized human oocytes in this study was approved by the relevant institutional ethics committees (LF MU 68/2022, RF 1/2015), and the UK’s National Research Ethics Service under the REC reference 11/EE/0346 (IRAS project ID 84952). Patients underwent ovarian stimulation for intra-cytoplasmic sperm injection (ICSI) as part of the treatment protocol. Only immature oocytes unsuitable for ICSI treatment were used in this study. All patients gave informed consent to donate their surplus oocytes for use in this study. Oocytes were collected and incubated in G-MOPS plus medium (Vitrolife, #10130) and G-IVF plus medium (Vitrolife, #10134) under paraffin oil (NidaCon, #NO-400K) at 37°C within 1 to 4 hours after ovarian retrieval, as previously described^67^. Oocytes were cultured in G-MOPS Plus or G-IVF Plus for a maximum of 24 to 48 hours until their maturation to metaphase II. Eggs were fixed 6 to 9 hours after polar body extrusion and stored in PBS supplemented with 0.02% sodium azide until preparation for immunofluorescence.

Human oocytes were vitrified and thawed using Kitazato Oocyte/Embryo Vitrification (#VT601) and Thawing Media (#VT602), respectively, following the manufacturer’s instructions^68^.

### Expression constructs and in vitro mRNA synthesis

To generate mRNA constructs for time-lapse imaging in oocytes, coding sequences of mSgo1 (transcript variant X1) and mSgo2 (transcript variant 1) were amplified from oocyte cDNA libraries generated using the SensiFAST cDNA synthesis kit (Bioline, #BIO-65053). Primers used for plasmid construction are listed in Table S1. First, pENTR/dTOPO-Sgo1, pENTR/dTOPO-Sgo1N61I, pENTR/dTOPO-hSgo1, pENTR/dTOPO-Sgo2, pENTR/dTOPO-H2B, pENTR/dTOPO-Map4-MTBD, were generated for subcloning into pGEMHE-mClover3-gateway destination vectors using primers P1-10 (Table S1) and Invitrogen™ pENTR™/TEV/D-TOPO™ Cloning Kit (Thermo Fisher Scientific, #K252520). Amplified sequences were subcloned into pGEMHE-mClover3 and pGEMHE-SNAP Gateway destination vectors using the Gateway™ LR Clonase™ II Enzyme Mix (Thermo Fisher Scientific, #11791020) to generate pGEMHE-mClover-mSgo1, pGEMHE-mClover-mSgo1N61I, pGEMHE-Sgo2-mClover, pGEMHE-hSgo1-EGFP, pGEMHE-SNAP-H2B, and pGEMHE-mClover3-MAP4-MTBD constructs. To generate the *hSGO1* (transcript variant A1) mRNA construct, vsv-Sgo1-EGFP plasmid was used (Addgene, #108494). Using restriction digestion with XhoI and NotI, the hSgo1 construct was subcloned into the pGEM-HE vector backbone for mRNA synthesis. All mRNAs were prepared using the HiScribe T7 ARCA mRNA Kit (NEB, # E2065S) according to the manufacturer’s instructions. Synthesized mRNAs were quantified using a Qubit RNA HS Assay Kit (Thermo Fisher Scientific, # Q32852).

The Mus musculus major satellite repeat (MajSat, V00846.1) targeting sequence, pGEM-T-MajSat, was a generous gift from Dr Maria-Elena Torres-Padilla^69^. Unlabeled *MajSat* sense and antisense RNA of the 234bp repeat of the mouse major satellite were transcribed using the MaxiScript kit (Ambion, #AM1312). For synthesizing the Cy3-labelled *MajSat* and *control* Cy3 RNA, the pGEM-T-MajSat plasmid with ScaI. For generating control RNA, the empty pGEM-T plasmid multiple cloning site was PCR amplified using primers 11,12 (Table S1). The linearized plasmid and amplified PCR product was used for performing in vitro transcription of the products using the T7 promoter. The HighYield T7 Cy3 RNA Labelling Kit (Jena Bioscience, #RNT-101-CY3) was used to label amplified sequences with Cy3-labelled UTP (UTP-X-Cy3, 35%), followed by DNA removal using Turbo™ DNAse (ThermoFisher). Purification of Cy3-uridine-labelled RNA was performed using Monarch® RNA Cleanup Kit (NEB, #T2040L).

### Microinjection of mouse and human oocytes and immature follicles

Oocytes and follicles were microinjected according to previously described protocols^70^. Oocytes and follicles were loaded onto an injection tray assembled by superimposing two coverslips separated by one (for mouse oocytes) and two (for mouse follicles and human oocytes) layers of 100 µm double-sided adhesive tape. For GV oocytes, 4-6 pl of the mRNAs were injected at the following concentrations *mSgo2-mClover3* at 65 ng/µl, *mClover3-Sgo1* at 65 ng/µl, *hSGO1-EGFP* at 20 or 80 ng/µl, *mMajSat-cy3* at 40 ng/µl, *Control RNA-Cy3* at 40 ng/µl, *SNAP-H2B* at 4 ng/µl, *mClover3-Map4-MTBD* at 80 ng/µl. Oocytes were allowed to express the mRNAs for 3 hours before dbcAMP was washed out to allow meiotic maturation.

Knockdown with short interfering RNAs (siRNAs) was performed by microinjecting siRNAs into follicle-enclosed oocytes isolated from 10-12-day old F1 females according to previously established protocols ^59^. The following siRNAs were used for Sgo1 knockdown. In this study, 2 different RNAi mixtures were used against MmSgol1 (NM_028232) from Qiagen (5’-CAGCAAATTGCTGTTGAAGAA-3’, SI01416674; 5’-CAGGAGAATTGCAGAGTACAA-3’, SI01416667). AllStars Negative Control (Qiagen, #1027281) was used as a control. Mouse follicles were microinjected with 6 pl of siRNAs at a needle concentration of 2 mM as described previously^59^.

### Immunofluorescence

To obtain mouse metaphase I oocytes and metaphase II eggs, GV oocytes were incubated in dbcAMP-free medium for ∼6 hours and ∼16 hours after release, respectively. Mouse oocytes and eggswere fixed for 30 minutes at 37°C in 100 mM HEPES (pH 7) (titrated with KOH), 50 mM EGTA (pH 7) (titrated with KOH), 10 mM MgSO4, 2% formaldehyde (MeOH-free), 0.2% Triton X-100, according to previously published methods^70,71^. Fixed oocytes were extracted with phosphate-buffered saline (PBS) with 0.5% Triton X-100 (PBT) overnight at 4°C and blocked in PBT with 5% BSA (PBT-BSA) for 6 hours at room temperature. All primary antibody incubations were performed overnight at 4°C in 5% PBT-BSA. After washing 3 times for 10 minutes with 5% PBT-BSA, oocytes were incubated with the secondary antibody mixture (20 µg/ml) for 1 hour at room temperature. Hoechst 33342 (Molecular Probes) was used at 100 µM for DNA staining with the final secondary antibody. Primary antibodies used for immunofluorescence analysis were as follows human anti-centromere (Antibodies Incorporated, #15-234-0001), rabbit anti-SGOL1 (gift from Dr. Yoshinori Watanabe)^58^, rat anti-α-tubulin (Santa Cruz, #sc-53030), rabbit anti-RNA pol II-pS2 (Abcam, #AB5095), rabbit anti-CENP-A (Cell Signaling, #2048), rabbit anti-SGOL2 (Biorbyt, #orb499806), rabbit anti-REC8 (produced by Cambridge Research Biochemicals using antigen protein corresponding to Mus musculus REC8 (aa25-286)), and rabbit anti-PP2A catalytic subunit (Merckmillipore, #05-421). Alexa Fluor 405, 488, 568 or 647 conjugated anti-human IgG, donkey IgG, mouse IgG, rabbit IgG or rat IgG (Thermo Fisher Scientific) were used as secondary antibodies.

Human oocytes were fixed for 30 minutes at 37°C in 100 mM HEPES (pH 7) (titrated with KOH), 50 mM EGTA (pH 7) (titrated with KOH), 10 mM MgSO4, 2% formaldehyde (MeOH-free), 0.2% Triton X-100, according to previously published methods^70^. Fixed oocytes were extracted with phosphate-buffered saline (PBS) with 0.5% Triton X-100 (PBT) and blocked in PBT with 5% BSA (PBT-BSA) overnight at 4°C. All primary antibody incubations were performed overnight at 4°C in 5% PBT-BSA. After washing 3 times for 10 minutes with 5% PBT-BSA, oocytes were incubated with the secondary antibody mixture (20 µg/ml) for 1 hour at room temperature. Hoechst 33342 (Molecular Probes, 100 µM) was used at for DNA staining with the final secondary antibody. Primary antibodies used for immunofluorescence analysis were as follows: human anti-centromere (Antibodies Incorporated, #15-234-0001), mouse anti-SGOL1 (Abnova, #HOO151648-M01), rabbit anti-SGO2 (Biotechne, #NBP1-83567), and rabbit anti-PP2A catalytic subunit (Cell Signaling, #2038S). Alexa Fluor 405, 488, 568, or 647 conjugated anti-human IgG, donkey IgG, rat IgG, mouse IgG, and rabbit IgG (Thermo Fisher Scientific) were used as secondary antibodies.

### Chromosome spreads

To obtain chromosome spreads of metaphase I and metaphase II chromosomes, oocytes were incubated in dbcAMP-free medium for ∼6 hours and ∼16 hours after release, respectively. To remove the zona, oocytes were washed briefly with 3-4 drops of Tyrode’s acid (Sigma, #T1788) followed by washing with M2 medium. Oocytes were swollen by incubation with a hypotonic solution (50% FBS, 50% ddH2O) on agarose gels (0.5% agarose in PBS) for 14 min at 37°C. Oocytes were fixed by dropping 12 µL of fixative (1% PFA, 0.15% Triton X-100, 3 mM DTT) into each well of a 15-well slide (MP Biomedicals, #096041505) in a humidified box. After overnight chromosome attachment to the slides, the slides were air dried and washed twice sequentially for 5 minutes each with spreading wash solution (0.08% Kodak Photoflo in 1L PBS, Laborimpex, #74257), PBS and immune wash solution (0.2% BSA, 0.1% Tween-20 in PBS). In case of CENPA antibody (cell signaling, #C51A7) lambda phosphatase treatment was performed before for 45 minutes at 30°C (3µL lambda Phosphatase, 30µL 10x PMP, 30µL MnCl2 stock, 237µl water). Spreads were blocked with 10% FBS, 2.5% BSA and 0.1% Tween-20 in PBS for 30 minutes at room temperature and antibody incubations were performed in PBT-BSA with primary antibodies or secondary antibodies for 1 hour at room temperature or overnight at 4°C. Hoechst 33342 (Molecular Probes) was used at 100 µM for DNA staining with the final secondary antibody. Slides were washed three times with a cleaning solution (0.01% Tween-20 in PBS), mounted with a 1:1 PBS-glycerol solution and sealed with nail varnish.

### Detection and inhibition of transcription in oocytes

The Click-iT™ RNA Alexa Fluor™ 488 Imaging Kit (ThermoFisher, #C10329) was used for detection of nascent transcripts in mouse oocytes. Isolated oocytes were transferred and washed in M2 (+dbcAMP) media supplemented with 5-ethynyluridine (EU, 1 mM) and incubated at 37°C for 3 hours. Oocytes were fixed as previously described, followed by extraction with PBT. The oocytes were then transferred to Click Reaction solution (172 μL Click Reaction Buffer, 8 μL CuSO4, 0.48 μL Alexa Fluo, 20 μL 1:10 diluted React Additive and 2 μL 10% TritonX-100) in the Terasaki plate and incubated for 30 minutes at room temperature. The oocytes were then washed with Click Rinse (Click Rinse solution diluted 1:100 with 10% TritonX-100) and PBS-Triton 0.1%. After two washes with each solution, oocytes were incubated in PBS-Triton 0.1% supplemented with Hoechst (100 µM) for 15-30 min in the dark before washing with PBS. The same procedure was used to check the efficacy of the transcription inhibitors triptolide and α-amanitin. To inhibit transcription in oocytes, Triptolide (Sigma, #T3652), and α-amanitin (Sigma, #A2263) were used at 25 µM for 3 hours and 20 µg/ml for 8 hours, respectively. Isolated oocytes were divided into control and treatment groups and then incubated with pre-warmed M2 media containing dbcAMP, inhibitor, SiR-DNA (Spirochrome, #SC007, stock 1 mM, working concentration 200 nM).

### Detection of satellite RNAs in oocytes

Detection of satellite transcripts was performed by RNA FISH using FISH probes against major satellite repeats according to protocols described before^72^. The probes used in this study are Cy3-O-GCGAGGAAAACTGAAAAAGG (MajSat antisense, PNAbio), FAM-O-TTGCCATATTCCACGTCC (MajSat sense, PNAbio). For metaphase I and II oocyte chromosome spreads, slides were washed twice briefly with 0.1% Tween-20 in RNase-free PBS followed by RNase-free PBS for 5 minutes at room temperature. For controls, slides were treated with RNase solution (100 µg/mL) for 30 minutes at 37°C and washed twice for 5 minutes with RNase-free PBS. PNA probes were used at 500 nM in freshly prepared hybridisation solution (60% formamide, 0.5% blocking reagent (Roche, #11096176001) in 20 mM Tris pH 7.4). Pre-warmed PNA probes were added to pre-warmed slides with chromosome spreads for 10 minutes at 85°C on a heating block. Slides were placed in a humidified chamber for 10 hours in the dark for probe hybridisation. Slides were sequentially washed in the dark with 50% (v/v) formamide in 2× SSC at 45°C for 15 minutes, 0.2× SSC at 63°C for 15 minutes, 2× SSC at 45°C for 5 minutes, and finally 2× SSC at room temperature for 5 minutes. For immunofluorescence, slides were washed in 1× PBS and blocked with 5% BSA (w/v) and 0.1% (v/v) Triton X-100 in 1× PBS for 1 hour at room temperature. Primary and secondary antibodies were incubated for 1 hour at room temperature. Slides were washed with RNase-free PBS before incubation with Hoechst solution for 1 hour at room temperature and washed three times with RNase-free PBS. 50% (v/v) glycerol-RNase-free PBS was used as mounting medium and sealed with coverslips (#1.5, VWR, 15165452) and nail varnish.

### Immunoblotting

For the detection of Sgo1 and RNA Pol II-pS2, 50 GV mouse oocytes were probed per lane. The harvested cells were washed briefly with PBS and equal numbers of oocytes from the control and experimental groups were snap frozen in liquid nitrogen. For oocyte lysis, cells were snap frozen and thawed a total of 3 times before mixing 5 µl NuPAGE™ LDS Sample Buffer (Thermo Fisher Scientific, #NP0007) supplemented with 100 mM DTT and 15 µl cell lysate and boiled at 95°C for 5 minutes. Protein samples, together with a pre-stained protein ladder (Thermo Fisher Scientific, #26619), were resolved on a 10-well NuPAGE 4-12% bis-tris protein gel, 1.0 mm thick (Thermo Fisher Scientific, # NP0322BOX) with 1X NuPAGE MOPS running buffer (Thermo Fisher Scientific, # NP0001). Proteins were transferred to a 0.45 µm PVDF membrane (Thermo Fisher Scientific, # LC2005) using 1X NuPAGE transfer buffer (Thermo Fisher Scientific, # NP0006) containing 10% methanol at 110 V for 1.5 hours. Blocking and antibody incubations were performed in PBS containing 5% skim milk and 0.05% Tween-20. The following primary antibodies were used: rabbit anti-SGOL1^58^ (1 µg/ml), rabbit anti-RNA Pol II-pS2 (Abcam, # AB5095, 2 µg/ml) and rabbit anti-DDB1 (Abcam, # ab109027, 0.04 µg/ml). Primary antibodies were incubated overnight at 4°C. Anti-rabbit HRP (Abcam, #ab205718) was used as secondary antibody and incubated for 1 hour at room temperature. Immunoblots were developed using Super Signal West Femto Maximum Sensitivity Substrate Femto (Thermo Fisher Scientific, #34094) and imaged using an Amersham Imager 600 (GE Healthcare).

### Staining of oocytes for live imaging

For time-lapse imaging of mouse oocytes, DNA was stained by incubating oocytes with M2 media containing 200 nM SiR-DNA (Spirochrome, #SC007, stock 1 mM, working concentration 200 nM). Time-lapse imaging of human oocytes were performed by incubating human oocytes with 5-SiR-Hoechst^73^ (stock concentration 1 mM, working concentration 50 nM). Dyes were reconstituted in DMSO.

### Confocal, super-resolution, and light-sheet microscopy

For confocal imaging, live mouse oocytes were imaged in pre-warmed M2 media and fixed oocytes in PBS with paraffin oil (NidaCon, #NO-400K) in a 35 mm glass bottom dish with a no. 1 coverslip (MatTek). Imaging with triptolide supplemented media was performed in Nunc™ Cell-Culture Treated Multidishes (Thermo Fisher, #176740) without oil. Confocal images were acquired using LSM 800/900/980 laser scanning confocal microscopes (Zeiss) equipped with an environmental incubator box and a 40× C-Apochromat 1.2 NA water immersion objective. Automatic 3D tracking using Autofocus screen was implemented for time-lapse imaging with a temporal resolution of 5 min^74,75^. For live imaging of mouse oocytes, images were acquired with an optical slice thickness of 2.5 µm confocal section with a z-stack interval of 2.0 µm covering approximately 40 µm. Typically, we recorded a volume centered around the chromosomes by 3D time-lapse imaging for multiple cells in parallel. mClover3, eGFP, and Alexa Fluor 488 was excited with a 488 nm laser line and detected at 493 – 571 nm. Alexa Fluor 568 was excited with a 561 nm laser line and detected at 571 – 638 nm. 5-SiR, SNAP-Cell 647-SiR, and Alexa Fluor 647 were excited with a 633 nm laser line and detected at 638 – 700 nm. Images were acquired under identical imaging conditions on the same microscope for the control and experimental groups.

Airyscan images were acquired using LSM800/900 confocal laser scanning microscopes equipped with an Airyscan module (Zeiss) and processed in ZEN (Zeiss) after acquisition. Care was taken to ensure that imaging conditions, including laser power, pixel dwell time and detector gain, did not cause phototoxicity, photobleaching or saturation.

Human oocytes were imaged in 2 ml of media under paraffin oil in a multi-well sample holder with four compartments (Viventis Microscopy Sàrl). Images were acquired with Viventis LS1 equipped with an environmental incubator box and 25X 1.1 NA water-dipping objective.

### Image analysis and quantification

Images were analysed in Fiji (https://imagej.nih.gov/ij/)^76^, Imaris (Bitplane), Arivis Vision4D or ZEN blue (ZEISS). The exported data were further processed in Microsoft Excel, R, and GraphPad Prism 9. In particular, a Gaussian filter was applied for noise reduction and maximum intensity projection of z-stacks for a subset of images (Zen, Zeiss). Airyscan processing was used when images were acquired using an Airyscan detector (Zen, Zeiss). Illustrations were created using Adobe Illustrator. Automated spot detection was used to identify kinetochore centers based on local maxima. For some cells, chromosome segmentation was performed in Imaris 9.3. Sister kinetochores were paired using filaments to generate seeds for individual chromosomes using scripts written in Matlab R2018b. Manual corrections were made where necessary. Inter-sister kinetochore distances were obtained by manual pairwise measurements between detected spots. PSSC was recorded when single chromatids with a single kinetochore were observed with no visible DNA signal in between two sister chromatids. Intensity measurements of RNA and proteins on chromosome spreads were performed using Fiji and Imaris (Bitplane). For centromeric proteins, automated spot detection was used to identify kinetochore centers based on local maxima and mean intensities were measured at the kinetochore. For proteins localized in an extended region around kinetochores, the region of interest was segmented in Fiji and a threshold was applied to mask the pericentromere. After thresholding, a binary mask was generated and multiplied by the original region of interest. The background was drawn manually and the mean background intensity was subtracted. Relative fluorescence intensity was calculated by normalization to the mean of the control cells. Quantification of REC8 and PP2A at the pericentromere in metaphase II eggs was performed using Arivis Vision4D. The machine learning segmenter was used to draw objects around the centromere and pericentromere and the background was drawn manually. After background subtraction, the mean intensity was normalized to the mean of control cells and relative intensity was plotted using GraphPad Prism 9. For quantification of Sgo1 levels in human oocytes, data were first processed with Huygens deconvolution for both Sgo1 and CREST channels to minimise noise. The same deconvolution parameters were used for both control and experimental data sets (maximum iteration 40, acuity 36.05). Total Sgo1 intensity on chromosomes was measured in Imaris after background subtraction and normalized to the mean of control cells.

### Statistical analysis

No statistical methods were used to predetermine sample size. Statistical analyses were performed with GraphPad v9.3.1 using unpaired two-tailed Student’s t-test (for absolute values) and two-tailed Fisher’s exact test (for categorical values). All box plots show median (horizontal black line), mean (small black squares), 25th and 75th percentiles (boxes), 10th and 90th percentiles (whiskers), percentiles (whiskers), and the 1st and 99th percentiles (circles). Standard deviation was used for error bars in box plots. P values are indicated with * sign (ns = not significant, *p≤0.05, **p≤0.01, ***p≤0.001, ****p≤0.0001). We tested the total number of intact sister chromatids against the total number of PSSC errors between two conditions, as indicated in each figure. Each experimental condition reported in this study includes at least two technical replicates, with majority of them consisting of having 3 or more experimental replicates. Donated human oocyte material is limited in quantity and therefore constrained sample size.

---

**Table S1:**
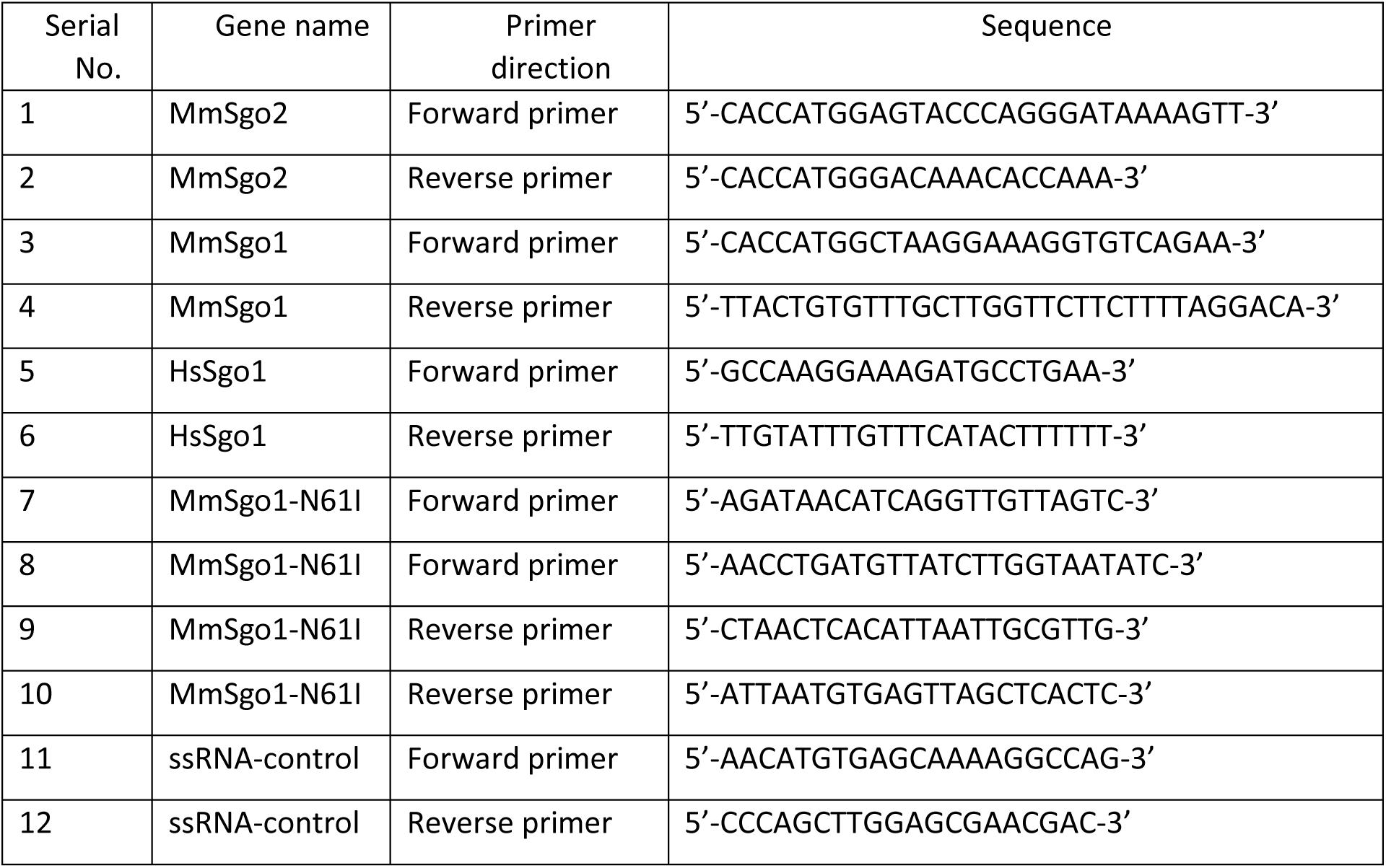
Oligonucleotides used in this study

**Table S2:** Details of donors of all human oocytes used in this study

| Figure panel | Donor Number | Age (years) | Mean age (years) | No. of GV oocyte (s) | No. of MI oocyte (s) | IVF clinic | Used in |
| --- | --- | --- | --- | --- | --- | --- | --- |
| Figures 6A and 6B | 1 | 26 | 35 | 1 | 0 | Bourn Hall Fertility Clinic | Immunofluorescence for measuring *SGO1* levels in metaphase I control oocytes |
|  | 2 | 28 |  | 3 | 0 |  |  |
|  | 3 | 39 |  | 1 | 0 |  |  |
|  | 4 | 38 |  | 1 | 0 |  |  |
|  | 5 | 35 |  | 0 | 5 |  |  |
|  | 6 | 33 |  | 1 | 0 |  |  |
|  | 7 | 39 |  | 0 | 2 |  |  |
|  | 8 | 39 |  | 1 | 0 |  |  |
|  | 9 | 36 |  | 1 | 0 |  |  |
|  | 10 | 37 |  | 0 | 2 |  |  |
|  | 11 | 38 |  | 1 | 0 |  |  |
|  | 12 | 41 |  | 1 | 0 |  |  |
|  | 13 | 34 |  | 1 | 0 |  |  |
|  | 14 | 32 |  | 0 | 1 |  |  |
|  | 15 | 30 |  | 0 | 2 |  |  |
|  | 16 | 28 |  | 1 | 0 |  |  |
|  | 17 | 34 |  | 0 | 1 |  |  |
|  | 18 | 38 |  | 0 | 1 |  |  |
|  | 19 | 32 |  | 1 | 0 |  |  |
|  | 20 | 39 |  | 1 | 0 |  |  |
|  | 21 | 43 |  | 0 | 2 |  |  |
|  | 22 | 29 |  | 1 | 0 |  |  |
| Figures 6C and 6D | 1 | 38 | 33 | 1 | 0 | Bourn Hall Fertility Clinic | Quantification of PSSC in metaphase II control eggs |
|  | 2 | 39 |  | 0 | 1 |  |  |
|  | 3 | 34 |  | 1 | 0 |  |  |
|  | 4 | 41 |  | 1 | 0 |  |  |
|  | 5 | 34 |  | 2 | 0 |  |  |
|  | 6 | 38 |  | 1 | 0 |  |  |
|  | 7 | 33 |  | 1 | 0 |  |  |
|  | 8 | 32 |  | 0 | 2 |  |  |
|  | 9 | 30 |  | 0 | 1 |  |  |
|  | 10 | 28 |  | 3 | 1 |  |  |
|  | 11 | 37 |  | 1 | 0 |  |  |
|  | 12 | 28 |  | 1 | 1 |  |  |
|  | 13 | 34 |  | 3 | 0 |  |  |
|  | 14 | 33 |  | 0 | 1 |  |  |
|  | 15 | 32 |  | 1 | 0 |  |  |
|  | 16 | 22 |  | 0 | 2 |  |  |
|  | 17 | 32 |  | 1 | 0 |  |  |
|  | 18 | 32 |  | 4 | 0 |  |  |
|  | 19 | 32 |  | 0 | 1 |  |  |
|  | 20 | 36 |  | 0 | 1 |  |  |
|  | 21 | 29 |  | 1 | 0 |  |  |
|  | 22 | 35 |  | 1 | 0 |  |  |
|  | 23 | 37 |  | 1 | 0 |  |  |
|  | 24 | 32 |  | 1 | 0 |  |  |
|  | 25 | 31 |  | 1 | 1 |  |  |
|  | 26 | 34 |  | 1 | 0 |  |  |
|  | 27 | 32 |  | 1 | 0 |  |  |
|  | 28 | 43 |  | 0 | 1 |  |  |
|  | 29 | 33 |  | 0 | 1 |  |  |
|  | 30 | 31 |  | 1 | 0 |  |  |
|  | 31 | 28 |  | 0 | 1 |  |  |
|  | 32 | 33 |  | 0 | 1 |  |  |
|  | 33 | 30 |  | 0 | 1 |  |  |
|  | 34 | 36 |  | 0 | 1 |  |  |
|  | 35 | 37 |  | 1 | 0 |  |  |
|  | 36 | 38 |  | 0 | 1 |  |  |
|  | 37 | 28 |  | 0 | 1 |  |  |
|  | 38 | 39 |  | 0 | 2 |  |  |
|  | 39 | 30 |  | 0 | 1 |  |  |
|  | 40 | 35 |  | 1 | 0 |  |  |
|  | 41 | 29 |  | 1 | 2 |  |  |
|  | 42 | 31 |  | 0 | 2 |  |  |
|  | 43 | 23 |  | 2 | 1 |  |  |
|  | 44 | 33 |  | 0 | 1 |  |  |
|  | 45 | 36 |  | 0 | 2 |  |  |
|  | 46 | 30 |  | 0 | 1 |  |  |
|  | 47 | 39 |  | 0 | 1 |  |  |
|  | 48 | 39 |  | 0 | 1 |  |  |
|  | 49 | 31 |  | 1 | 1 |  |  |
|  | 50 | 33 |  | 0 | 2 |  |  |
|  | 51 | 33 |  | 0 | 1 |  |  |
|  | 52 | 40 |  | 0 | 1 |  |  |
|  | 53 | 42 |  | 0 | 1 | Reprofit<br>Internationa<br>I Clinic |  |
|  | 54 | 31 |  | 0 | 2 |  |  |
| Figures<br>6E–J | 1 | 38 | 33 | 1 | 0 | Bourn Hall<br>Fertility<br>Clinic | Quantification of<br>PSSC in control eggs<br>(overall) |
|  | 2 | 39 |  | 0 | 1 |  |  |
|  | 3 | 34 |  | 1 | 0 |  |  |
|  | 4 | 41 |  | 1 | 0 |  |  |
|  | 5 | 34 |  | 2 | 0 |  |  |
|  | 6 | 38 |  | 1 | 0 |  |  |
|  | 7 | 33 |  | 1 | 0 |  |  |
|  | 8 | 32 |  | 0 | 2 |  |  |
|  | 9 | 30 |  | 0 | 1 |  |  |
|  | 10 | 28 |  | 3 | 1 |  |  |
|  | 11 | 37 |  | 1 | 0 |  |  |
|  | 12 | 28 |  | 1 | 1 |  |  |
|  | 13 | 34 |  | 3 | 0 |  |  |
|  | 14 | 33 |  | 0 | 1 |  |  |
|  | 15 | 32 |  | 1 | 0 |  |  |
|  | 16 | 22 |  | 0 | 2 |  |  |
|  | 17 | 32 |  | 1 | 0 |  |  |
|  | 18 | 32 |  | 4 | 0 |  |  |
|  | 19 | 32 |  | 0 | 1 |  |  |
|  | 20 | 36 |  | 0 | 1 |  |  |
|  | 21 | 29 |  | 1 | 0 |  |  |
|  | 22 | 35 |  | 1 | 0 |  |  |
|  | 23 | 37 |  | 1 | 0 |  |  |
|  | 24 | 32 |  | 1 | 0 |  |  |
|  | 25 | 31 |  | 1 | 1 |  |  |
|  | 26 | 34 |  | 1 | 0 |  |  |
|  | 27 | 32 |  | 1 | 0 |  |  |
|  | 28 | 43 |  | 0 | 1 |  |  |
|  | 29 | 33 |  | 0 | 1 |  |  |
|  | 30 | 31 |  | 1 | 0 |  |  |
|  | 31 | 28 |  | 0 | 1 |  |  |
|  | 32 | 33 |  | 0 | 1 |  |  |
|  | 33 | 30 |  | 0 | 1 |  |  |
|  | 34 | 36 |  | 0 | 1 |  |  |
|  | 35 | 37 |  | 1 | 0 |  |  |
|  | 36 | 38 |  | 0 | 1 |  |  |
|  | 37 | 28 |  | 0 | 1 |  |  |
|  | 38 | 39 |  | 0 | 2 |  |  |
|  | 39 | 30 |  | 0 | 1 |  |  |
|  | 40 | 35 |  | 1 | 0 |  |  |
|  | 41 | 29 |  | 1 | 2 |  |  |
|  | 42 | 31 |  | 0 | 2 |  |  |
|  | 43 | 23 |  | 2 | 1 |  |  |
|  | 44 | 33 |  | 0 | 1 |  |  |
|  | 45 | 36 |  | 0 | 2 |  |  |
|  | 46 | 30 |  | 0 | 1 |  |  |
|  | 47 | 39 |  | 0 | 1 |  |  |
|  | 48 | 39 |  | 0 | 1 |  |  |
|  | 49 | 31 |  | 1 | 1 |  |  |
|  | 50 | 33 |  | 0 | 2 |  |  |
|  | 51 | 33 |  | 0 | 1 |  |  |
|  | 52 | 40 |  | 0 | 1 |  |  |
|  | 53 | 42 |  | 0 | 1 |  |  |
|  | 54 | 31 |  | 0 | 2 |  |  |
|  | 55 | 38 |  | 1 | 0 |  |  |
|  | 56 | 39 |  | 0 | 1 |  |  |
|  | 57 | 34 |  | 1 | 0 |  |  |
|  | 58 | 41 |  | 1 | 0 |  |  |
|  | 59 | 34 |  | 2 | 0 |  |  |
|  | 60 | 38 |  | 1 | 0 |  |  |
|  | 61 | 33 |  | 1 | 0 |  |  |
|  | 62 | 32 |  | 0 | 2 |  |  |
|  | 63 | 30 |  | 0 | 1 |  |  |
|  | 64 | 28 |  | 3 | 1 |  |  |
|  | 65 | 37 |  | 1 | 0 |  |  |
|  | 66 | 28 |  | 1 | 1 |  |  |
|  | 67 | 34 |  | 3 | 0 |  |  |
|  | 68 | 33 |  | 0 | 1 |  |  |
|  | 69 | 32 |  | 1 | 0 |  |  |
|  | 70 | 22 |  | 0 | 2 |  |  |
|  | 71 | 32 |  | 1 | 0 |  |  |
|  | 72 | 32 |  | 4 | 0 |  |  |
|  | 73 | 32 |  | 0 | 1 |  |  |
|  | 74 | 28 | 31 | 3 | 0 | Bourn Hall<br>Fertility<br>Clinic | Quantification of<br>PSSC in eggs<br>microinjected with |
|  | 75 | 28 |  | 1 | 0 |  |  |
|  | 76 | 34 |  | 1 | 0 |  |  |
|  | 77 | 37 |  | 2 | 0 |  |  |
| | 78 | 28 | | 1 | 0 | | SGO1 mRNA (20 or 80 ng/μl) |
|  | 79 | 22 |  | 1 | 0 |  |  |
|  | 80 | 38 |  | 1 | 0 |  |  |
|  | 81 | 32 |  | 2 | 0 |  |  |
|  | 82 | 29 |  | 1 | 2 |  |  |
|  | 83 | 30 |  | 1 | 0 |  |  |
|  | 84 | 35 |  | 1 | 1 |  |  |
|  | 85 | 37 |  | 2 | 1 |  |  |
|  | 86 | 32 |  | 2 | 0 |  |  |
|  | 87 | 31 |  | 2 | 1 |  |  |
|  | 88 | 34 |  | 1 | 0 |  |  |
|  | 89 | 28 |  | 1 | 1 |  |  |
|  | 90 | 31 |  | 1 | 0 |  |  |
|  | 91 | 30 |  | 0 | 2 |  |  |
|  | 92 | 35 |  | 1 | 0 |  |  |
|  | 93 | 30 |  | 1 | 0 |  |  |
|  | 94 | 23 |  | 3 | 2 |  |  |
| Figure 6K | 1 | 28 | 30 | 3 | 1 | Bourn Hall Fertility Clinic | Quantification of PSSC in eggs from the same donor (control) |
|  | 2 | 28 |  | 1 | 1 |  |  |
|  | 3 | 34 |  | 3 | 0 |  |  |
|  | 4 | 22 |  | 0 | 2 |  |  |
|  | 5 | 32 |  | 1 | 0 |  |  |
|  | 6 | 29 |  | 1 | 0 |  |  |
|  | 7 | 35 |  | 1 | 0 |  |  |
|  | 8 | 37 |  | 1 | 0 |  |  |
|  | 9 | 32 |  | 1 | 0 |  |  |
|  | 10 | 31 |  | 1 | 1 |  |  |
|  | 11 | 34 |  | 1 | 0 |  |  |
|  | 12 | 28 |  | 0 | 1 |  |  |
|  | 13 | 30 |  | 0 | 1 |  |  |
|  | 14 | 23 |  | 2 | 1 |  |  |
| | 15 | 28 | 30 | 3 | 0 | Bourn Hall Fertility Clinic | Quantification of PSSC in eggs from the same donor (eggs treated with 20 or 80 ng/μl SGO1 mRNA) |
|  | 16 | 28 |  | 1 | 0 |  |  |
|  | 17 | 34 |  | 1 | 0 |  |  |
|  | 18 | 22 |  | 1 | 0 |  |  |
|  | 19 | 32 |  | 2 | 0 |  |  |
|  | 20 | 29 |  | 1 | 2 |  |  |
|  | 21 | 35 |  | 1 | 1 |  |  |
|  | 22 | 37 |  | 2 | 1 |  |  |
|  | 23 | 32 |  | 2 | 0 |  |  |
|  | 24 | 31 |  | 2 | 1 |  |  |
|  | 25 | 34 |  | 1 | 0 |  |  |
|  | 26 | 28 |  | 1 | 1 |  |  |
|  | 27 | 30 |  | 0 | 2 |  |  |
|  | 28 | 23 |  | 3 | 2 |  |  |
| Figures<br>S5E–G | 1 | 36 | 33 | 0 | 1 | Bourn Hall<br>Fertility<br>Clinic | Quantification of<br>PSSC in control eggs |
|  | 2 | 29 |  | 1 | 0 |  |  |
|  | 3 | 35 |  | 1 | 0 |  |  |
|  | 4 | 37 |  | 1 | 0 |  |  |
|  | 5 | 32 |  | 1 | 0 |  |  |
|  | 6 | 31 |  | 1 | 1 |  |  |
|  | 7 | 34 |  | 1 | 0 |  |  |
|  | 8 | 32 |  | 1 | 0 |  |  |
|  | 9 | 43 |  | 0 | 1 |  |  |
|  | 10 | 33 |  | 0 | 1 |  |  |
|  | 11 | 31 |  | 1 | 0 |  |  |
|  | 12 | 28 |  | 0 | 1 |  |  |
|  | 13 | 33 |  | 0 | 1 |  |  |
|  | 14 | 30 |  | 0 | 1 |  |  |
|  | 15 | 36 |  | 0 | 1 |  |  |
|  | 16 | 37 |  | 1 | 0 |  |  |
|  | 17 | 38 |  | 0 | 1 |  |  |
|  | 18 | 28 |  | 0 | 1 |  |  |
|  | 19 | 39 |  | 0 | 2 |  |  |
|  | 20 | 30 |  | 0 | 1 |  |  |
|  | 21 | 35 |  | 1 | 0 |  |  |
|  | 22 | 29 |  | 1 | 2 |  |  |
|  | 23 | 31 |  | 0 | 2 |  |  |
|  | 24 | 23 |  | 2 | 1 |  |  |
|  | 25 | 33 |  | 0 | 1 |  |  |
|  | 26 | 36 |  | 0 | 2 |  |  |
|  | 27 | 30 |  | 0 | 1 |  |  |
|  | 28 | 39 |  | 0 | 1 |  |  |
|  | 29 | 39 |  | 0 | 1 |  |  |
|  | 30 | 31 |  | 1 | 1 |  |  |
|  | 31 | 33 |  | 0 | 2 |  |  |
|  | 32 | 33 |  | 0 | 1 |  |  |
|  | 33 | 29 | 31 | 1 | 2 |  | Quantification of<br>PSSC in eggs treated |
|  | 34 | 30 |  | 1 | 0 |  |  |
| | 35 | 35 | | 1 | 1 | Bourn Hall Fertility Clinic | with *SGO1* mRNA (20 ng/μl) |
|  | 36 | 37 |  | 2 | 1 |  |  |
|  | 37 | 32 |  | 2 | 0 |  |  |
|  | 38 | 31 |  | 2 | 1 |  |  |
|  | 39 | 34 |  | 1 | 0 |  |  |
|  | 40 | 28 |  | 1 | 1 |  |  |
|  | 41 | 31 |  | 1 | 0 |  |  |
|  | 42 | 30 |  | 0 | 2 |  |  |
|  | 43 | 35 |  | 1 | 0 |  |  |
|  | 44 | 30 |  | 1 | 0 |  |  |
|  | 45 | 23 |  | 3 | 2 |  |  |
| Figures S5H–J | 1 | 38 | 33 | 1 | 0 | Bourn Hall Fertility Clinic | Quantification of PSSC in control eggs |
|  | 2 | 39 |  | 0 | 1 |  |  |
|  | 3 | 34 |  | 1 | 0 |  |  |
|  | 4 | 41 |  | 1 | 0 |  |  |
|  | 5 | 34 |  | 2 | 0 |  |  |
|  | 6 | 38 |  | 1 | 0 |  |  |
|  | 7 | 33 |  | 1 | 0 |  |  |
|  | 8 | 32 |  | 0 | 2 |  |  |
|  | 9 | 30 |  | 0 | 1 |  |  |
|  | 10 | 28 |  | 3 | 1 |  |  |
|  | 11 | 37 |  | 1 | 0 |  |  |
|  | 12 | 28 |  | 1 | 1 |  |  |
|  | 13 | 34 |  | 3 | 0 |  |  |
|  | 14 | 33 |  | 0 | 1 |  |  |
|  | 15 | 32 |  | 1 | 0 |  |  |
|  | 16 | 22 |  | 0 | 2 |  |  |
|  | 17 | 32 |  | 1 | 0 |  |  |
|  | 18 | 32 |  | 4 | 0 |  |  |
|  | 19 | 32 |  | 0 | 1 |  |  |
| | 20 | 28 | 32 | 3 | 0 | Bourn Hall Fertility Clinic | Quantification of PSSC in eggs treated with *SGO1* mRNA (80 ng/μl) |
|  | 21 | 28 |  | 1 | 0 |  |  |
|  | 22 | 34 |  | 1 | 0 |  |  |
|  | 23 | 37 |  | 2 | 0 |  |  |
|  | 24 | 37 |  | 1 | 0 |  |  |
|  | 25 | 28 |  | 1 | 0 |  |  |
|  | 26 | 22 |  | 1 | 0 |  |  |
|  | 27 | 38 |  | 2 | 0 |  |  |

